# Two DNA binding domains of Mga act in combination to suppress ectopic activation of meiosis-related genes in mouse embryonic stem cells

**DOI:** 10.1101/2020.07.21.215079

**Authors:** Kousuke Uranishi, Masataka Hirasaki, Yuka Kitamura, Yosuke Mizuno, Masazumi Nishimoto, Ayumu Suzuki, Akihiko Okuda

## Abstract

Mouse embryonic stem cells (ESCs) have high potential for meiotic entry, like germ cells. Although the physiological meaning of this potential is not known, it is certain that a rigid safeguarding system is required to prevent ectopic onset of meiosis. PRC1.6, a non-canonical PRC1, is known for its suppression of precocious and ectopic meiotic onset in germ cells and ESCs, respectively, in which MGA has important roles in DNA binding as well as in constructing the complex as a scaffolding component. As a salient feature, MGA bears two distinct DNA-binding domains termed bHLHZ and T-box. However, how these features contribute to the functions of PRC1.6, particularly in the repression of meiotic genes, remains largely obscure. Here, we demonstrated that both DNA binding domains of Mga repress distinct sets of genes in murine ESCs, and substantial numbers of meiosis-related genes are included in both gene sets. In addition, our data demonstrated that both DNA binding domains of Mga, in particular bHLHZ, are crucially involved in repressing the expression of *Meiosin*, which plays essential roles in meiotic entry in collaboration with *Stra8*, revealing at least part of the molecular mechanisms that link negative and positive regulation of meiotic onset.

## INTRODUCTION

Like Myc family proteins, MGA bears a basic helix-loop-helix leucine zipper (bHLHZ) domain and dimerizes with MAX, which also has a bHLHZ domain for its binding to specific DNA sequences (E-box) (Hurlin et al., 1999). However, unlike Myc family proteins, MGA belongs to the MXD/MAD transcriptional repressor family, which functions antagonistically against Myc activity by competing with Myc for dimerization with MAX as well as for subsequent DNA binding to E-box sequences (Baudino and Cleveland, 2001; Grandori et al., 2000; Llabata et al., 2020; Schaub et al., 2018). In addition to the common bHLHZ domain, MGA also bears a MAX-independent DNA binding domain called the T-box domain, which directs binding to a specific DNA sequence, the T-box motif (Baudino and Cleveland, 2001; Grandori et al., 2000; Hurlin et al., 1999).

Although MGA by itself is able to repress transcription from genes with a T-box-containing promoter and can repress transcription from genes with an E-box-containing promoter as well when dimerized with MAX, MGA/MAX heterodimer is known to be incorporated into one particular type of polycomb repressive complex 1 (PRC1), whereby PRC1.6 is the scaffolding component of the complex (Gao et al., 2012; Stielow et al., 2018). PRC 1 constitutes a family of six distinct subtypes, PRC1.1 to PRC1.6, which differ in their subunit composition (Gao et al., 2012; Hauri et al., 2016; Scelfo et al., 2019). The PRC1 family is largely classified into two groups of canonical (PRC1.2 and PRC1.4) and non-canonical (PRC1.1, PRC1.3, PRC1.5 and PRC1.6) proteins (Gao et al., 2012; Morey et al., 2013; Tavares et al., 2012; Zepeda-Martinez et al., 2020). Even among non-canonical PRC1 members, distinct as well as common roles appeared to be assigned to each complex, such as a specific role for PRC1.3 and PRC1.5 in X chromosome inactivation(Almeida et al., 2017). We and others have previously demonstrated that PRC1.6 is involved in repressing the transcription of meiosis-related genes in ESCs and germ cells as a rather unique role of this complex (Endoh et al., 2017; Maeda et al., 2013; Stielow et al., 2018; Suzuki et al., 2016). Although the physiological importance of the potential of ESCs for meiotic onset is totally obscure, a rigid system that masks this potential is absolutely required for safeguarding against ectopic onset of meiosis that would lead to not only loss of pluripotency but also extensive cell death (Suzuki et al., 2016). Therefore, it is important to know how PRC1.6 exerts its role by repressing the transcription of meiosis-related genes in ESCs. As a biochemically unique feature among PRC1s, PRC1.6 bears sequence-specific DNA binding proteins such as MGA and E2F6 that are directly involved in the binding of PRC1.6 to the genome in a DNA-sequence specific manner (Gao et al., 2012; Stielow et al., 2018). However, how PRC1.6 uses these special characteristics, particularly in the regulation of meiosis-related genes, have not been intensively explored thus far.

Here, we scrutinized the roles of two DNA binding domains of Mga in murine ESCs. Our data revealed that genes subjected to negative regulation by these two DNA binding domains are rather distinct from each other. However, a substantial number of meiosis-related genes were included in both gene sets. Our data also revealed that bHLHZ-dependent regulation of meiosis-related genes almost exclusively reflected the role of Mga within PRC1.6, while T-box-dependent regulation of Mga was relatively less tightly linked to PRC1.6. Moreover, our data identified a molecular link between the positive and negative regulation of meiotic entry in which meiotic entry promoted by the retinoic acid (RA)–*Stra8* axis (Anderson et al., 2008; Baltus et al., 2006; Dokshin et al., 2013; Mark et al., 2008) is further potentiated by de-repression of *Meiosin* gene, which encodes an indispensable partner for Stra8 (Ishiguro et al., 2020) upon impairment of the transcriptional repressing activity of PRC1.6 that is known to occur at the onset of meiosis in germ cells (Suzuki et al., 2016).

## MATERIALS AND METHODS

### Cell culture

Mouse ESCs (EBRTcH3) (Masui et al., 2005) and their derivatives were cultured as described previously (Suzuki et al., 2016). HEK293FT cells were cultured with Dulbecco’s modified Eagle’s medium containing 10% fetal bovine serum. All-trans retinoic acid (RA) was used at 100 nmol/ml for activating *Stra8* and other meiosis-related genes.

### Genetic manipulation of *Mga* in ESCs by lentivirus-mediated CRISPR-Cas9 system

Oligonucleotides listed in Table S1 were used to edit exons 2 and 20 of *Mga* individually. pLentiCRISPRv2 (#52961; Addgene) carrying specific oligonucleotides and psPAX2 (#52961; Addgene) and pLP-VSVG (Invitrogen) vectors were co-transfected into HEK293 cells. Lentiviruses recovered from transfected cells were used to infect parental ESCs with polybrene (8 μg/ml). Then, the infected ESCs were subjected to puromycin selection (1 μg/ml) for 6 days. Subsequently, the resultant puromycin-resistant ESC colonies were recovered individually, and their genomic DNA was used to identify ESC clones that had been subjected to appropriate modifications in *Mga* loci.

### DNA microarray and Gene Ontology (GO) analyses

DNA microarray analyses were conducted as described previously (Hirasaki et al., 2018). Gene Ontology (GO) analyses were performed using DAVID web tools (http://david.abcc.ncifcrf.gov).

### RNA isolation, reverse transcription, and quantitative PCR (qPCR)

Total RNA from parental ESCs or their derivatives was used to obtain cDNA by reverse transcription. The cDNA was then used for qPCR using TaqMan probes or by SYBR Green-based methods. Oligonucleotides used for SYBR Green-based methods and TaqMan probes are listed in Table S1 and Table S2, respectively. All samples were tested in triplicate and the results were normalized with *Gapdh* expression levels.

### Chromatin immunoprecipitation (ChIP) analyses

Chromatin immunoprecipitation analyses were conducted using parental ESCs and two different *Mga*-mutant ESC lines (2×10^6^ cells each), as described previously (Suzuki et al., 2016). Genomic DNA purified from immunoprecipitated chromatins were used for SYBR Green-based qPCR. Specific primers used in these analyses are listed in Table S1.

### ChIP-seq analyses

ChIP-seq analyses were performed as described previously (Hirasaki et al., 2018). The raw data of sequence reads were subjected to filtration for quality checks (FASTX Toolkit 0.0.13, **http://hannonlab.cshl.edu/fastx_toolkit/index.html** fastq_quality_filter, parameters -Q 33 -q 20 -p 75) and subsequently mapped to the mouse genome (mm10 assembly) using bowtie2-2.2.5 software. The resulting SAM files were converted to BAM format with the aid of SAMtools (version 0.1.19) and then the obtained BAM files were subjected to peak calling using MACS2.1.1 software with the parameters of -q 1e5 -c for each IgG file. Peaks identified by the procedure were subjected to the two nearest genes association rules (within 1 kb) of the Genomic Regions Enrichment of Annotations Tool (http://bejerano.stanford.edu/great/public/html/splash.php). BigWig files were generated by subtracting signal values obtained with input samples using bamCompare software from deepTools 2.0. MultiBigwigSummary (a tool of Deeptools) was used to compute the average scores for each 5 kb genomic region with a 1-kb bin size. For genomic binding sites of *Pcgf6* and *Mga* in ESCs, publicly available ChIP-seq data (Stielow et al., 2018) deposited in the ArrayExpress database with the accession number of E-MATB-6007 were used. A *de novo* motif search was performed using the MEME-ChIP v5.1.1 website (http://meme-suite.org/tools/meme-chip) (Machanick and Bailey, 2011). A genomic region with a 5-kb length that carries a canonical transcription start site (TSS) of each objective gene at the center was used for the analyses. E-box, T-box, and E2F-binding sites yielded from the search were originated from reports by Arttu et al., Xu et al., and Fornes et al., respectively (Fornes et al., 2020; Jolma et al., 2013; Xu et al., 2007).

### Immunostaining and alkaline phosphatase staining

Immunocytochemical analyses were conducted as described previously (Suzuki et al., 2016) using wild-type and two different *Mga*-mutant ESC lines that had been treated or untreated with RA. Alkaline phosphatase staining was performed using a Leukocyte Alkaline Phosphatase kit (Sigma) according to the manufacturer’s protocol.

### Histone extraction

Histone solutions were obtained with wild-type and two different *Mga*-mutant ESC lines according to Stein and Mitchell (1988).

### Western blot analysis

Nuclear extracts and histone extraction solutions from parental and two *Mga*-mutant ESC lines were used for western blot analyses of Mga and histones, respectively, as described previously (Suzuki et al., 2016). Antibody against human MGA that cross-reacts with mouse Mga was kindly provided by Dr. Stielow et al.(Stielow et al., 2018) at Philipps-University of Marburg in Germany, while the other primary antibodies used are listed in Table S2.

## ACCESSION NUMBERS

DNA microarray and ChIP-seq data were deposited in the Gene Expression Omnibus (GEO) and Sequence Read Archives (SRA), respectively, under accession numbers of GSE154073 and PRJNA647006.

## RESULTS

### Generation of ESCs producing Mga mutants that lack either the bHLHZ or T-box domain

To assess the roles of the two DNA binding domains of Mga in ESCs, we conducted genetic manipulation to produce mutant Mga proteins lacking either T-box (ΔT ESCs) or bHLHZ (ΔbHLHZ ESCs), which are encoded in the second and 20th exons of the *Mga* gene, respectively, using the CRISPR-Cas9 system (Fig S1A). Then, screening procedures such as RT-PCR were performed to identify appropriately mutated clones (Fig. S1B–S1E).

Since Mga is known to control expression from a large number of genes as a potent transcriptional repressor, we first examined whether the levels of histone modifications were globally affected by the deletion of either DNA binding domain. However, no noticeable changes in the levels of H2AK119ub, H3K4me^3^, H3K27me^3^, and H3K27Ac were evident in either ΔT or ΔbHLHZ ESCs (Fig S2A). We also found that the levels of alkaline phosphatase activity (Fig. S2B) and the expression of pluripotent and naïve genes that serve as indicators of the undifferentiated state of ESCs (Fig. S2C, D) were comparable among wild-type and mutant ESC lines, indicating that, unlike ESCs that are completely null for *Mga* (Washkowitz et al., 2015), ΔT and ΔbHLHZ ESCs fairly retained their undifferentiated state.

### bHLHZ and T-box independently regulate their target genes

To compare the global expression profile between ΔT and ΔbHLHZ ESCs unbiasedly, we conducted DNA microarray analyses (Fig. S3A, B). Although Venn diagrams showed that 36 and 11 genes were commonly up- and downregulated between the two mutant ESC lines, a much larger number of genes were identified as having expression levels that were altered exclusively in either ESC mutant (Fig. 1A, Table S3, S4), indicating that both DNA binding domains of Mga target rather distinct sets of genes from each other. These data also indicated that the two DNA binding domains of Mga act in combination to control the expression levels of many genes. We also compared genes that were elevated more than 2-fold between ΔT and ΔbHLHZ ESCs by constructing a two-dimensional scatter plot. Notably, this comparison led to the finding that, with only two exceptions, the vast majority of profoundly activated genes (more than 5-fold) were rather restrictively activated in either ΔT or ΔbHLHZ ESCs (Fig. 1B), further strengthening the notion that both DNA binding domains control rather distinct targets to each other. Next, we conducted a GO classification to correlate gene expression changes with overall molecular functions. First, these analyses indicated that meiosis-related genes are rather prominently activated in both ΔbHLHZ and ΔT ESCs (Fig. 1C, D). Contrary to the activated genes, neither analyses of downregulated genes in ΔbHLHZ nor those in ΔT ESCs yielded a term whose P-value was significantly small (Fig. 1C, D), indicating that these gene sets represent rather disordered assemblies in which the biological roles of individual genes are rather diversified from one another. These data also supported the notion that major role of Mga is not the activation but the repression of transcription. Because of these reasons, we focused on activated, but not downregulated, gene sets in either *Mga*-mutant ESCs for subsequent analyses.

**Figure 1.**
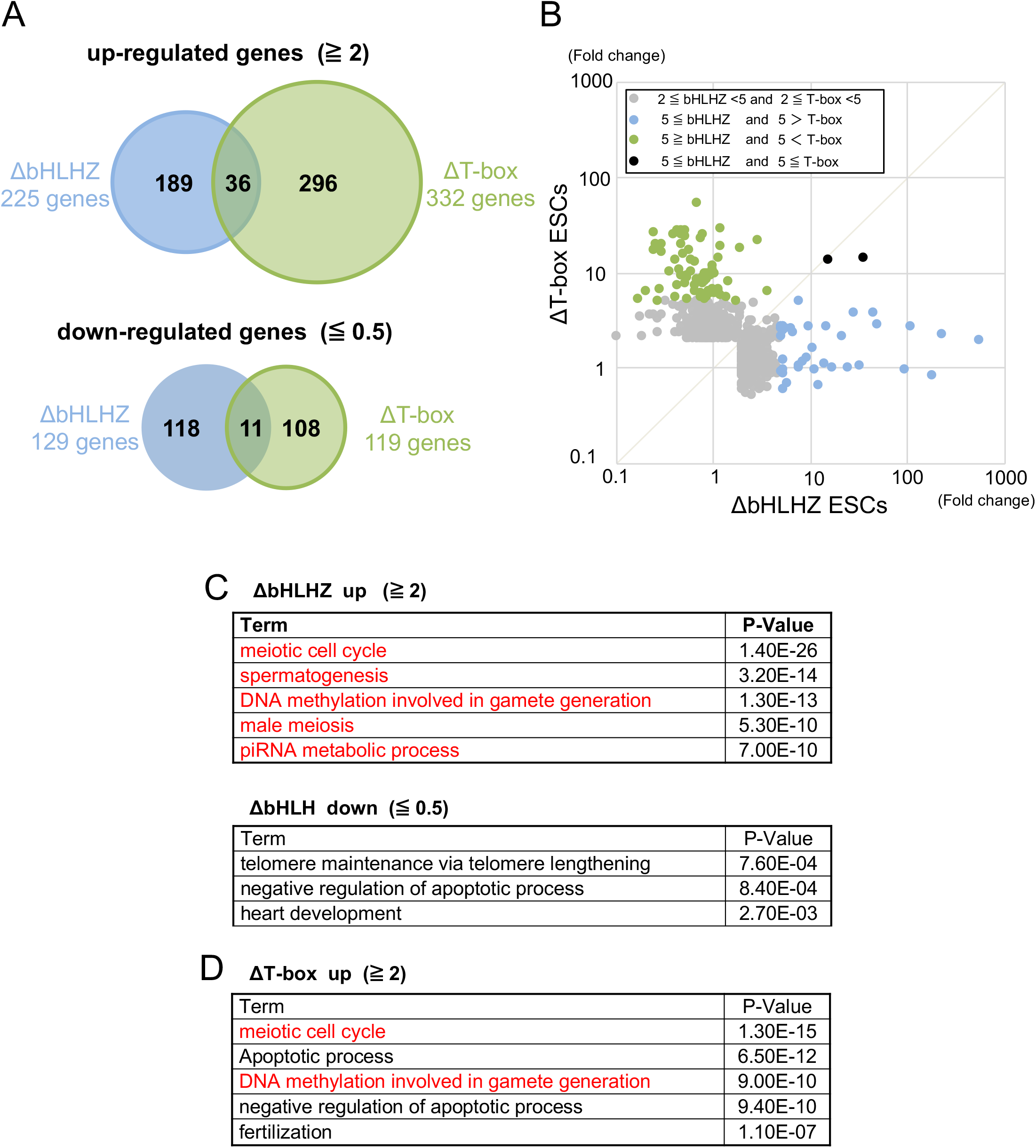
Comparison of alterations in global expression profiles between ΔbHLHZ and ΔT ESCs. (A) Venn diagrams showing comparisons of genes that increased (upper panel) or decreased (lower panel) more than two-fold in their signal values from DNA microarray analyses between ΔbHLHZ and ΔT ESCs. (B) Comparison of genes that were upregulated by more than two-fold between ΔbHLHZ and ΔT ESCs using a scatter plot. (C) GO analyses of genes showing more than 2-fold up- (upper panel)- or down- (lower panel)-regulation in ΔbHLHZ ESCs. Top five GO terms in terms of small *P*-values are shown in each analysis. However, terms whose P-values were larger than 10^−2^ were removed from the lists. (D) GO analyses of genes showing more than 2-fold up- (upper panel)- or downregulation in ΔT ESCs. Data are shown as in C in which no term that met the criteria for listing was obtained from the analyses of downregulated genes in ΔT ESCs.

### Cross-reference of activated genes in *Mga*-mutant ESCs with *de novo* binding site for Pcgf6

Since the MGA/MAX heterodimer is known to be incorporated in PRC1.6, we explored how tightly *Mga* mutation-dependent derepression is linked to impairment of PRC1.6. To this end, we cross-referenced the genes upregulated in ΔbHLHZ (225 genes) or ΔT ESCs (332 genes) with publicly available ChIP sequence data of genomic binding sites for Pcgf6, which is the specific component of PRC1.6 in ESCs (Endoh et al., 2017; Gao et al., 2012; Scelfo et al., 2019; Stielow et al., 2018). These analyses revealed that only approximately 36% and 21% of genes activated in ΔbHLHZ and ΔT ESCs, respectively, overlapped with the publicly reported genomic binding sites for Pcgf6 (Fig. 2A), indicating that substantial portions of upregulated genes in these ESC mutants are not subjected to the PRC1.6-dependent regulation. Next, we conducted a *de novo* motif search of DNA sequences around Pcgf6-binding genomic sites of these overlapping genes. These analyses allowed confirm that E-box and T-box sequences were relatively enriched in genes that are activated in ΔbHLHZ and ΔT ESCs, respectively, albeit T-box sequence enrichment in ΔT ESCs was much less significant compared with prominent E-box enrichment within the activated genes in ΔbHLHZ ESCs (Fig. 2B). We also noted that the activated genes in ΔbHLHZ ESCs and, even more so, those activated in ΔT ESCs, enriched E2F6 motif in their Pcgf6-binding sites (Fig. 2B), implicating that E2F6/DP1(or DP2) heterodimer is also involved in recruiting PRC1.6 to their gene promoters. Next, we investigated the possibility that the magnitude of induction of the activated genes in either *Mga*-mutant ESC line is influenced by the presence of *de novo* binding sites for Pcgf6. Our data demonstrated that, among the activated genes in ΔbHLHZ ESCs, genes with *de novo* Pcgf6-binding sites in general showed much more pronounced activation than genes that do not bear the site (Fig. 2C), implicating that the former gene group is generally much more tightly repressed by the HLHZ domain of Mga in ESCs and this difference is probably explained by the difference in the involvement of PRC1.6 in their transcriptional regulation. However, unexpectedly, no such a bias was evident among activated genes in ΔT ESCs (Fig. 2C). We will discuss the possibilities that may account for these data later (see Discussion).

**Figure 2.**
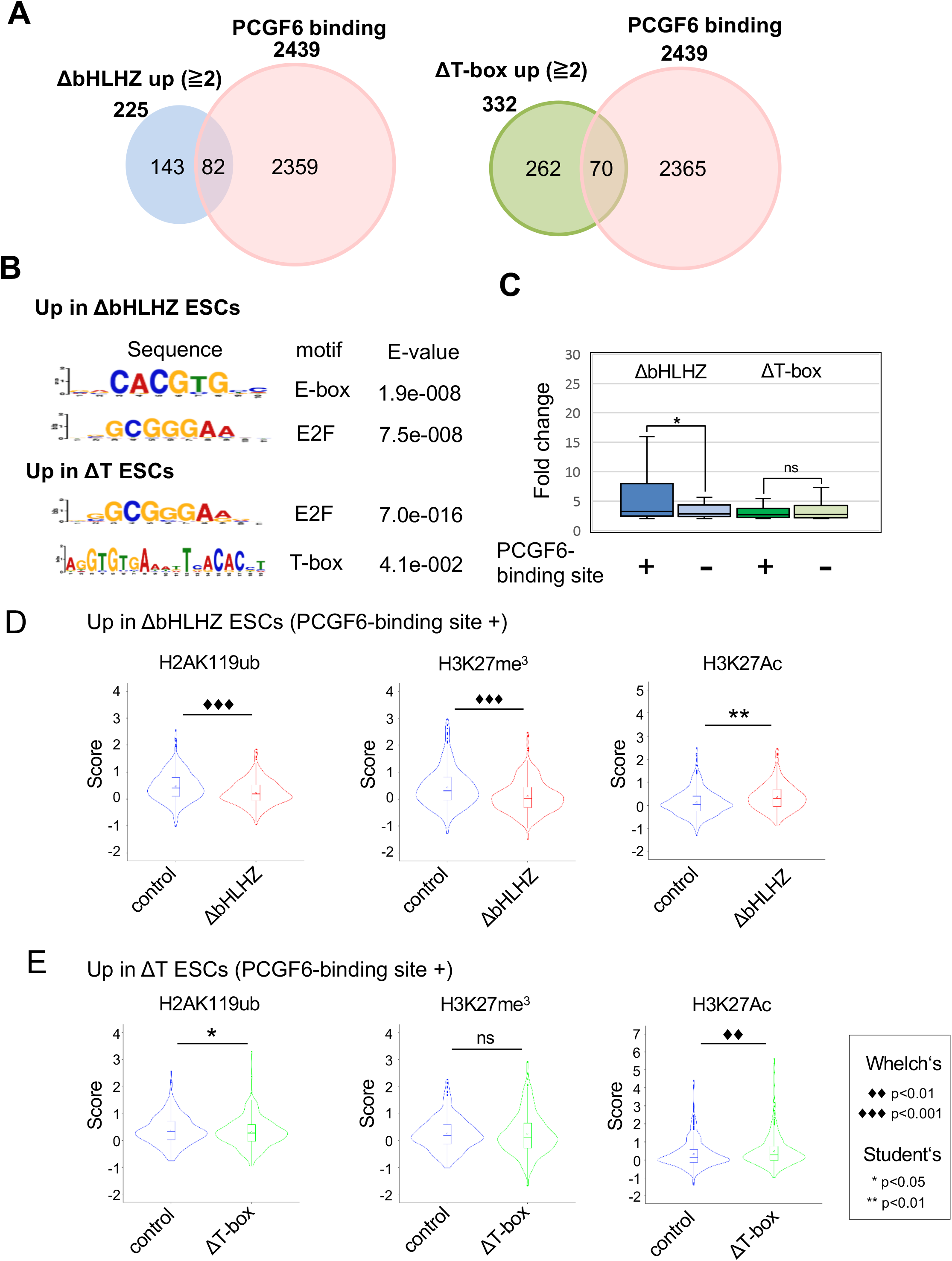
Cross-reference of the activated genes in *Mga*-mutant ESCs with publicly reported Pcgf6-binding sites. (A) Venn diagram showing the relationship between genes activated in either ΔbHLHZ or ΔT ESCs and publicly reported Pcgf6 binding sites. (B) A *de novo* sequence motif search. Pcgf6-binding regions of genes activated in either ΔbHLHZ or ΔT ESCs were individually used to search for prevalent motifs unbiasedly using MEME-ChIP software(Machanick and Bailey, 2011) and then data for E-box, T-box, and E2F motif sequences were picked up and shown. Data are shown with statistical significance (E-value). (C) Examination of the effect of the presence of Pcgf6-binding sites on the magnitude of derepression in *Mga*-mutant ESCs. Genes activated in ΔbHLHZ or ΔT ESCs were individually divided into two groups according to the presence or absence of publicly reported genomic Pcgf6-binding sites as in A. Fold-activation in the respective groups indicated is displayed with box plots. *P<0.05; ns: not significant (D, E) Examination of epigenetic alterations in genomic Pcgf6-binding site-containing genes that are activated in ΔbHLHZ or ΔT ESCs. Scores of ChIP-seq data for H2AK119ub (left), H3K27me3 (middle), and H3K27KAc (right) of Pcgf6-binding site-containing activated genes in ΔbHLHZ (D) and those in ΔT ESCs (E) in parental and correspondent mutant ESCs are displayed in violin plots. For statistical significance examination, F-test values were first obtained for the respective data. Then, Student’s t-test was conducted in cases in which the F-test value was larger than 0.05, while Welch’s t-test was conducted when its value was smaller than 0.05.

Next, we performed ChIP-experiments followed by high-throughput sequencing (ChIP-seq) to inquire whether histone modification levels of genes activated in either ΔbHLHZ or ΔT ESCs were influenced by the presence or absence of de *novo* binding sites for Pcgf6. First, to confirm the validity of our ChIP-seq data, we inspected the actual profiles of histone modifications across *Mov1011* and *Tdrkh* that appeared to be prominently activated specifically in ΔbHLHZ and ΔT ESCs, respectively (Table S3). Consistent with the alteration in their expression profiles, the levels of H3K27Ac were increased around the *Mga*-binding regions of *Mov10l1* and *Tdrkh* concomitantly with the deletion of the bHLHZ and T-box domains in ESCs, respectively, while the levels of histone modifications of H2A K119ub and H3K27me^3^ were inversely correlated with H3K27Ac levels (Fig. S4A). We also conducted locus-specific ChIP-qPCR analyses focused on the *Mga*-binding sites of these two genes and obtained consistent results (Fig. S4B), further validating the ChIP-seq data. Then, we used these data for more global analyses. Specifically, we examined whether pronounced activation of genes in ΔbHLHZ ESCs because of the presence of a publicly reported genomic binding site for Pcgf6 was accompanied with expected alterations in their histone modification levels. These analyses revealed that the genes with a genomic binding site for Pcgf6 showed an obvious tendency of decreased levels of H2AK119ub and H3K27me3 specifically in ΔbHLHZ ESCs that was coupled to a concomitant increase in their H3K27Ac levels (Fig. 2D). However, the levels of histone modifications of H2AK119ub and H3K27me3 were only marginally declined among genes that do not bear *de novo* Pcgf6 binding sites, and no statistically significant difference in the levels of histone H3K27Ac was evident from the comparison with those from parental ESCs (Fig. S5A). We also conducted analyses of genes activated specifically in ΔT ESCs. However, no statistically significant data were obtained from these analyses except for marginal decreases and increases in the modification levels of H2AK119ub and H3K27Ac among the genes with *de novo* binding sites for Pcgf6, respectively (Fig. 3E and Fig. S5B).

**Figure 3.**
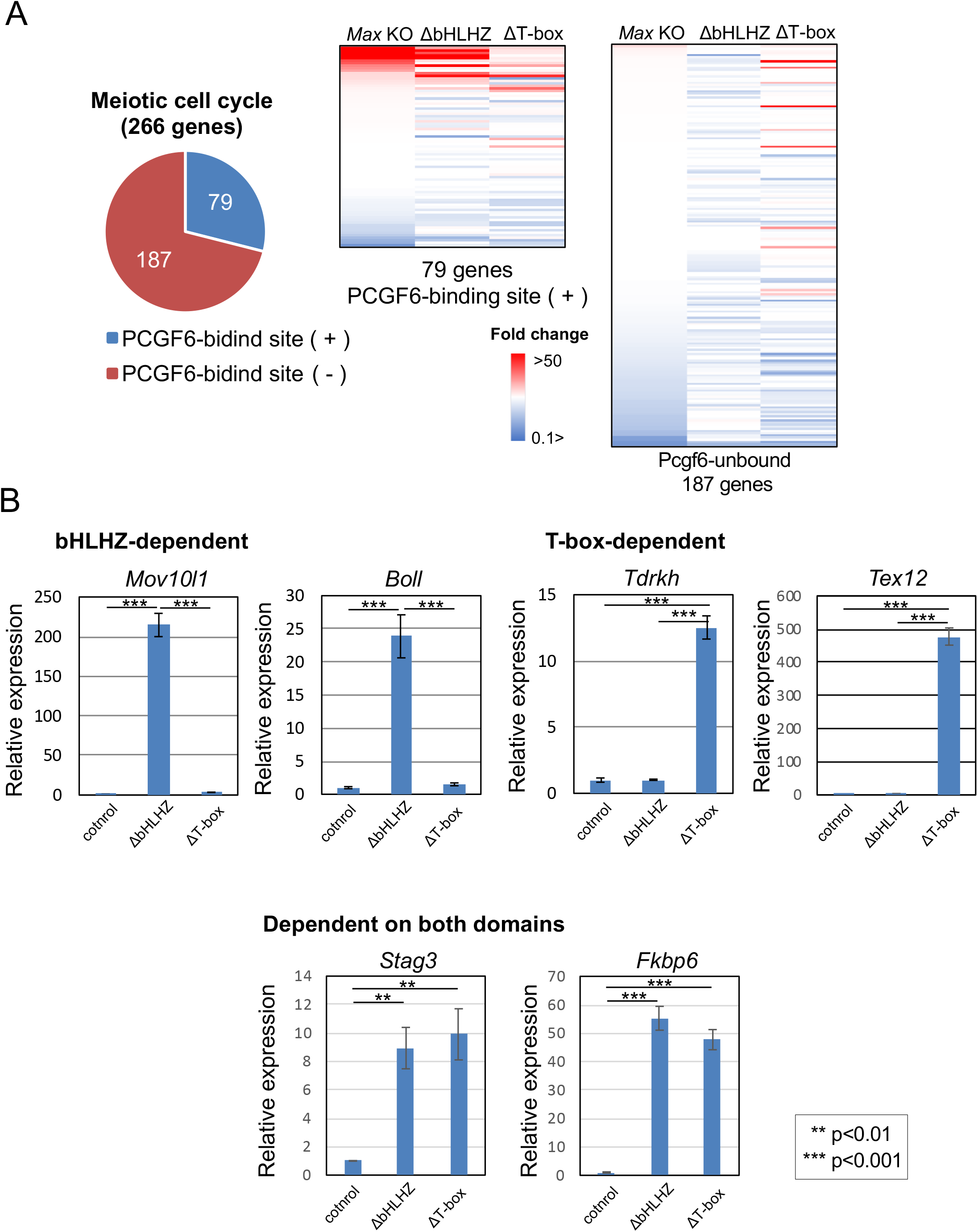
Alterations in the expression of meiosis-related genes in ΔbHLHZ and ΔT ESCs. (A) Heat map showing the comparison of alterations in the expression levels of meiosis-related genes (GO:0051321) among *Max*-KO and two *Mga*-mutant (ΔbHLHZ and ΔT) ESCs. First, meiosis-related genes were classified into two subsets according to the presence or absence of *de novo* genomic binding sites for Pcgf6, as shown in a pie chart. Then, data from these two subsets were individually used for generating heat maps. (B) qPCR analyses of representative genes activated in ΔbHLHZ and/or ΔT ESCs. Two genes were selected from each group of genes activated in either ΔbHLHZ (*Mov10l1, Boll*) or ΔT ESCs (*Tdrkh, Tex12*) or genes activated in both *Mga*-mutant ESCs (*Stag3, Fkbp6*) for the analyses. Data from parental ESCs were arbitrarily set to one. Data represents mean ± standard deviation of three independent experiments. Student’s t-test was conducted to examine statistical significance.

### Similarities in the activation profiles of meiosis-related genes between *Max*-KO and ΔbHLHZ ESCs

GO analyses indicated that, like *Max*-KO ESCs, both ΔbHLHZ and ΔT *Mga*-mutant ESCs also activated meiosis-related genes significantly. Thus, we decided to compare the activation profiles of those genes among these three mutant ESCs. Before conducting the comparisons, meiosis-related genes were classified into two groups according to the presence or absence of *de novo* binding sites for Pcgf6. Then, DNA microarray data of individual meiotic cell cycle genes from *Max*-KO, ΔbHLHZ, and ΔT ESCs were plotted with a heat map using data from parental ESCs as references. First, these analyses revealed that *Max*-KO ESCs rather restrictively activated a subset of meiosis-related genes that bear *de novo* binding sites for Pcgf6 (Fig. 3A), supporting the notion that *Max* expression ablation-mediated activation of meiosis-related genes reflects the consequence of impairment of the function of PRC1.6. We also noted that ΔbHLHZ ESCs were relatively more similar to *Max*-KO ESCs than to ΔT ESCs in the alterations of transcriptional profiles, especially the activated genes in this gene set (Fig. 3A). These results are rather to be expected, since bHLHZ domain of MGA is closely linked to MAX functionally by forming an MGA/MAX heterodimer through their common bHLHZ domains. Although ΔT ESCs also activated a substantial number of genes classified into this gene set, these were largely distinct from the activated genes in *Max*-KO ESCs and ΔbHLHZ ESCs. The analyses of meiosis-related genes that lack *de novo* binding sites for Pcgf6 revealed that, although some of them were rather uniquely activated in ΔT ESCs, none of those genes showed an obvious tendency of upregulation in either *Max*-KO or ΔbHLHZ ESCs. Next, we performed quantitative PCR (qPCR) to validate the DNA microarray data. These analyses revealed that the expression profiles that were consistent with the DNA microarray data were obtained from all of six meiosis-related genes that we selected as genes activated exclusively in either ΔbHLHZ (*Mov10l1* and *Boll*) or ΔT ESCs (*Tdrkh* and *Tex12*) or activated in both *Mga*-mutant (*Stag3* and *Fkbp6*) ESC lines (Fig. 3B).

### Marginal activation of two cell-specific genes in ESCs by deletion of the T-box of Mga

We have previously demonstrated that ablation of *Max* expression was accompanied by conspicuous induction of two cell-specific genes including *Zscan4*, as well as meiosis-related genes in ESCs. Therefore, we decided to examine whether two cell-specific genes are also activated in ΔbHLHZ and/or ΔT ESCs. First, we used DNA microarray data to inspect the expression profile of two cell-specific genes. Since 18 genes among the two cell-specific genes (Mikkelsen et al., 2007) are also categorized as meiosis-related genes, we classified the two cell-specific gene set into two subsets according to their relationship with meiosis. Then, these two subsets were further classified according to the presence or absence of *de novo* genomic binding sites for Pcgf6, as shown in Figure 3A. These analyses revealed that *Max*-KO ESCs rather restrictively activated genes that bear *de novo* Pcgf6-binding sites among two cell-specific genes with meiosis-related characteristics (Fig. 4A, upper left panel), implying that these genes are under the control of PRC1.6. However, the vast majority of the remaining two cellspecific genes that were activated in *Max*-KO ESCs lacked *de novo* Pcgf6-binding sites (Fig. 4A, lower left panel), indicating that the profound activation of two cell-specific genes with no meiosis-related characteristics in *Max*-KO ESCs was not attributed to the disruption of the function of PRC1.6. These analyses also revealed that there was a significant similarity in the activation profile of the former subset between *Max*-KO ESCs and ΔbHLHZ ESCs, whereas the activation profile of this gene set in ΔT ESCs was rather distinct (Fig. 4A, upper left panels). These analyses also revealed that the other gene set with no meiosis-related characteristics showed no tendency of activation in ΔbHLHZ ESCs, irrespective of the presence or absence of *de novo* Pcgf6-binding sites, whereas those without *de novo* Pcgf6-binding sites, but not those with them, showed marginal similarity in the activation profile between *Max*-KO and ΔT ESCs (Fig. 4A, lower left and right panels). *Dux*, a master regulator of two cell-specific genes, has been shown to be subjected to negative regulation by PRC1.6 (Cossec et al., 2018). However, *Dux* is not included as one of the above two cell-specific genes, since this gene was not identified as a master regulator for two cell-specific genes at the time of classification (Mikkelsen et al., 2007). Therefore, we conducted qPCR for determining whether Mga mutations were accompanied by alterations in the expression levels of *Dux* and *Zscan4*, the latter a well-known direct target of *Dux* (Cossec et al., 2018; De Iaco et al., 2017; Hendrickson et al., 2017). These analyses revealed that an obvious activation of these genes was not observed in ΔbHLHZ ESCs, but instead ΔT ESCs showed moderately high expression of these genes (Fig. 4B). Taken together, these results indicated that some representative two cellspecific genes including *Dux* were activated in ΔT ESCs, but that was not sufficient for the rather broad activation of two cell-specific genes.

**Figure 4.**
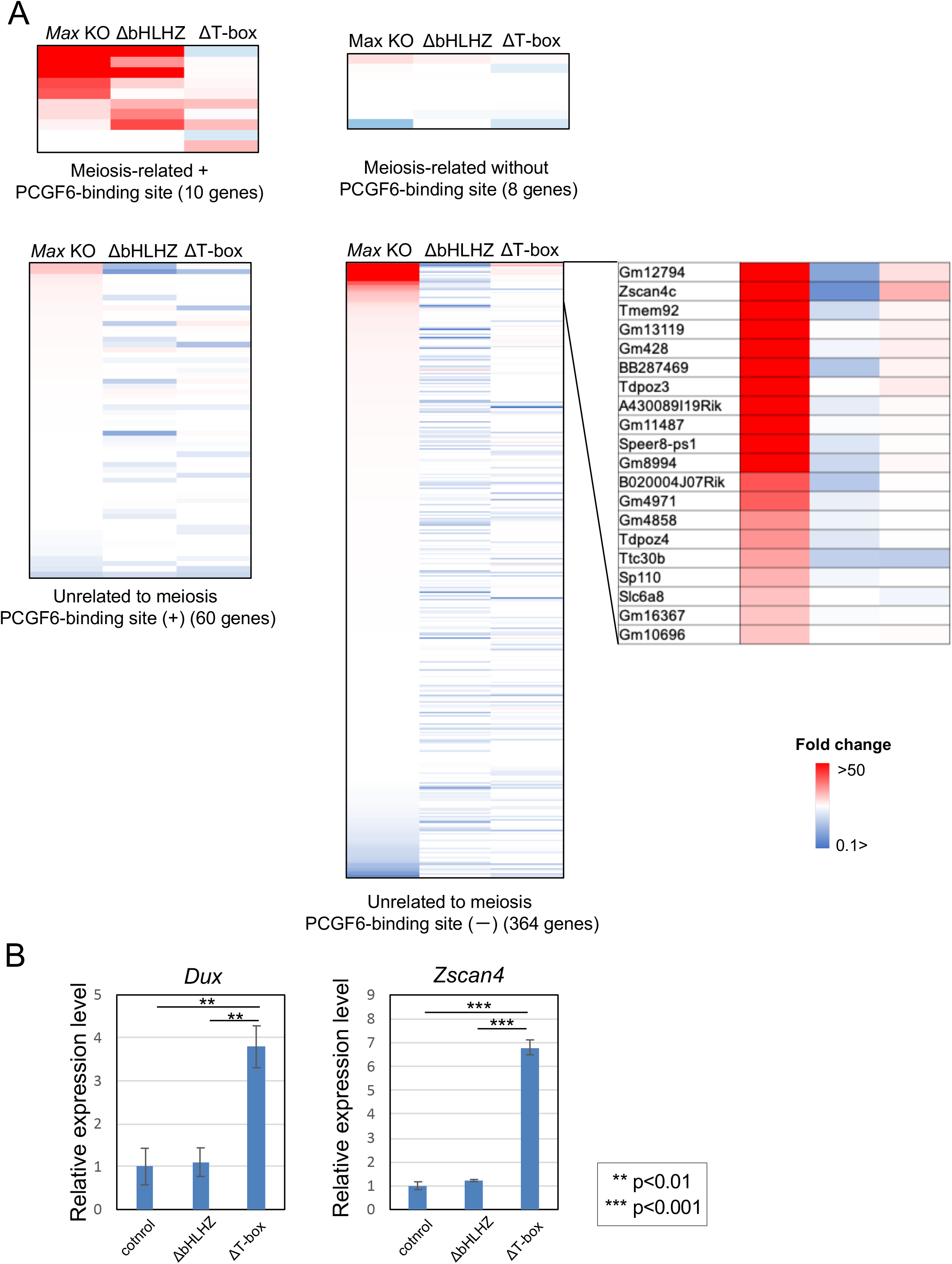
Comparison of the expression profiles of two cell-specific genes in ΔbHLHZ and ΔT ESCs with that of *Max*-KO ESCs. (A) Heat map showing the comparison of alterations in the expression levels of two cell-specific genes among *Max*-KO and two *Mga*-mutant ESC lines. Two cell-specific gene sets (442 genes) were classified into two subgroups in which one subtype (18 genes) was composed of genes that were also categorized into meiotic cell cycle genes that constitute the GO term (GO:0051321) and the other subset (424 genes) was composed of the remaining two cellspecific genes with no meiosis-related characteristics. The DNA microarray data of these two gene sets were further classified according to the presence or absence of *de novo* genomic binding sites for Pcgf6 and these four subsets were individually used to generate heat maps. The data were lined-up according to the order of fold-induction in *Max*-KO ES cells. Among genes classified into the subset with neither meiosis-related characteristics nor *de novo* Pcgf6-binding sites, the top 20 genes in terms of the magnitude of activation in *Max*-KO ESCs were highlighted by enlarging an upper portion of the heat map and are shown on the right. (B) qPCR analyses of the expression of representative two cell-specific genes. Expression levels of *Dux* and *Zscan4* genes in ΔbHLHZ and ΔT ESCs were quantified by qPCR. Data from parental ESCs were arbitrarily set to one. Data represents mean ± standard deviation of three independent experiments. Student’s t-test was conducted to examine statistical significance.

### Deletion of the bHLHZ domain of Mga sensitizes ESCs to their response to retinoic acid-mediated induction of meiotic entry

While PRC1.6 is assigned to negative regulation against meiotic onset, RA is known as a molecule that strongly promotes meiotic onset (Bowles et al., 2006; Koubova et al., 2006) and STRA8 is the central player in this signaling cascade (Anderson et al., 2008; Baltus et al., 2006; Dokshin et al., 2013; Mark et al., 2008; Zhou et al., 2008). In this context, one can assume that these two negative and positive regulators do not function independently, but there are certain molecular mechanisms that adjust the balance of the relative dominance between these mutually opposing regulations appropriately to smoothen the onset of meiosis. However, all previous reports including ours indicate that *Stra8* is not a direct target of the negative regulation of PRC1.6 (Endoh et al., 2017; Suzuki et al., 2016). Interestingly, however, our data indicated that *Meiosin*, which has recently been shown to encode an indispensable partner for Stra8 (Ishiguro et al., 2020), is a direct downstream target of PRC1.6 whose expression is elevated profoundly and marginally in ΔbHLHZ and ΔT ESCs, respectively (see Table S3). Histone modification analyses by ChIP-seq as well as locus-specific ChIP-qPCR strongly supports the notion that *Meiosin*, but not *Stra8*, was subjected to Mga (bHLHZ domain)-dependent regulation of PRC1.6 (Fig. S6A, B). Moreover, since the STRA8/MEIOSIN complex has been shown to bind to their own gene promoters to construct a feedforward regulatory loop (Ishiguro et al., 2020), we examined whether RA-dependent induction of *Stra8* was further potentiated in ΔbHLHZ ESCs in which the expression of *Meiosin* was significantly elevated. In accordance with this hypothesis, we found that ΔbHLHZ ESCs showed much more pronounced induction of *Stra8* gene expression than wild-type and ΔT ESCs upon RA treatment (Fig. 5A). We also examined the expression levels of several target genes of the Stra8/Meiosin complex including *Meiosin* itself. These analyses revealed that, although RA treatment by itself did not result in appreciable induction, genes upregulated in ΔbHLHZ ESCs (*Meiosin, Sycp3, Taf7l, Slc25a31, Dazl*), but not those upregulated in ΔT ESCs (*Tdlrkh, Hormad2l*), were further activated by treatment with RA (Fig. 5B). We consider that the difference in responsiveness against RA between ΔbHLHZ and ΔT ESCs can be explained by the differences in the expression levels of *Meiosin* in which only marginal activation of *Meiosin* is evident in ΔT ESC mutant. We also found that high expression of Stra8 and Sycp3 in RA-treated ΔbHLHZ ESCs could be confirmed by immunocytochemistry (Fig. 5C).

**Figure 5.**
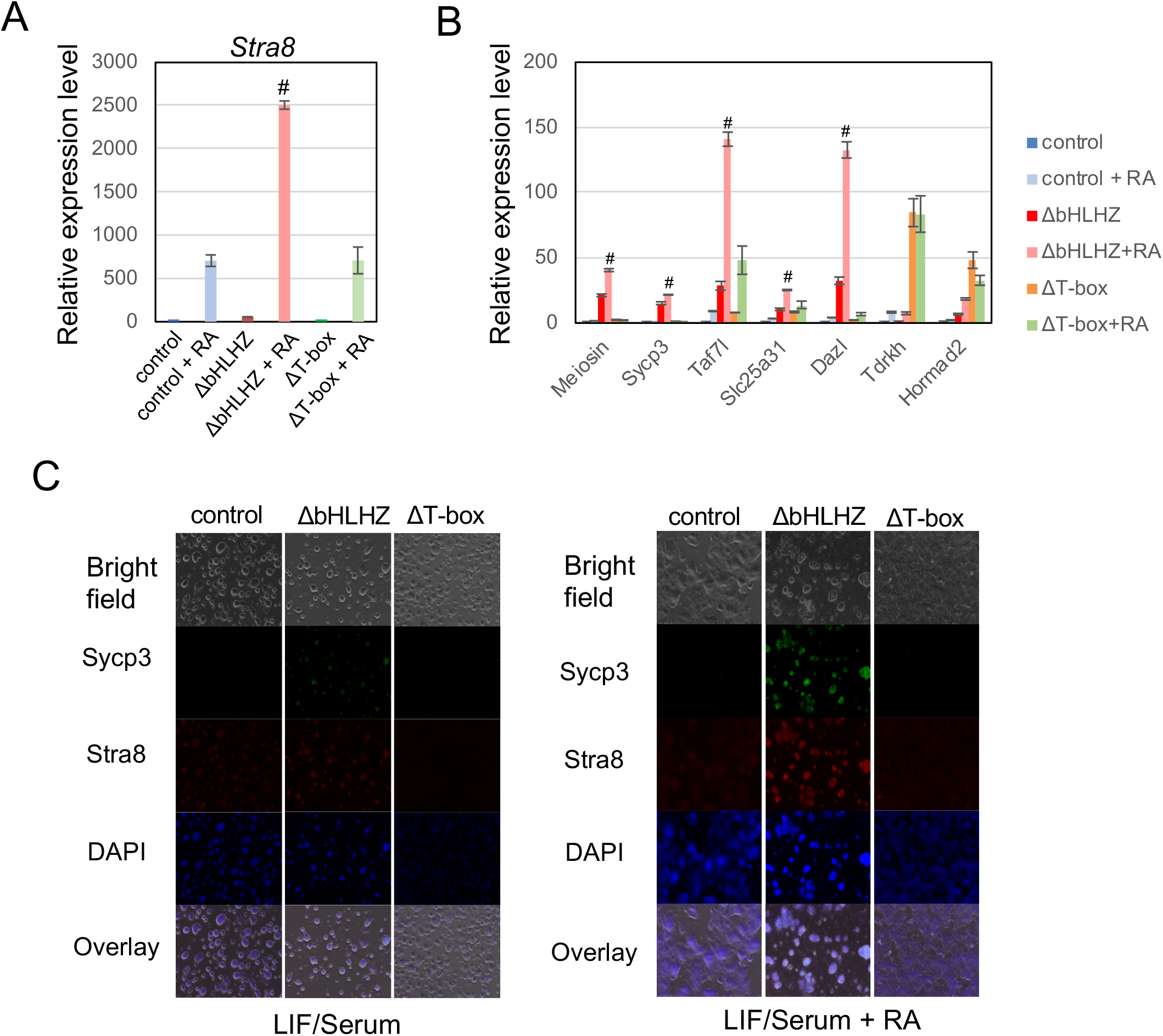
Meiosin could serve as a critical molecule that determines the relative dominance of positive and negative regulation of meiosis. (A) Strong potentiation in RA-mediated induction of *Stra8* in ΔbHLHZ ESCs. qPCR analyses were performed to determine *Stra8* expression levels in ΔbHLHZ, ΔT, and parental ESCs that were treated or untreated with RA for 2 days. Expression levels of *Stra8* in untreated parental ESCs were arbitrarily set to one. Data represents mean ± standard deviation of three independent experiments. Student’s t-test was conducted to examine statistical significance. Values that were statistically different (*P*<0.05) from all other five values obtained in ESCs that were distinct in *Mga* loci and/or RA-treatment are indicated by #. (B) Meiosis-related genes activated in ΔbHLHZ ESCs, but not those activated in ΔT ESCs, were further activated by RA-treatment. qPCR quantification of the expression levels of meiosis-related genes that were subjected to bHLHZ-dependent (*Meiosin, Sycp3, Taf7l, Slc25a31, Dazl*) and T-box dependent (*Tdrkh, Hormad2*) regulation in parental and *Mga*-mutant (ΔbHLHZ and ΔT) ESCs cultured with or without RA. The expression level of each gene in untreated parental ESCs was arbitrarily set to one. Data represents mean ± standard deviation of three independent experiments. Data were subjected to statistical analyses as in A. (C) Immunocytochemical analyses of parental and *Mga*-mutant (ΔbHLHZ and ΔT) ESCs that were treated with or without RA. Indicated ESCs were treated or untreated with RA for 2 days and subjected to co-immunostaining with antibodies against Sycp3 and Stra8.

## DISCUSSION

We and others have previously demonstrated that disruption of PRC1.6 leads to profound activation of meiosis-related genes in ESCs as well as in germ cells, but only marginal activation is evident in somatic cells such as fibroblasts (Endoh et al., 2017; Maeda et al., 2013; Stielow et al., 2018; Suzuki et al., 2016). However, a recent report demonstrated that Pcgf6-deficient somatic cells also showed rather high expression of meiosis-related genes (Liu et al., 2020), contrasting with data from other reports. It is known that transcriptional repression of meiosis-related genes by PRC1.6 in pluripotent early embryonic cells is taken over by DNMT3B-mediated DNA methylation from the beginning of their differentiation (Auclair et al., 2014; Borgel et al., 2010; Greenberg and Bourc’his, 2019). Therefore, given that repression by PRC1.6 is an absolutely required prerequisite step for DNMT3B-mediated silencing, we assume that differences in the activated state of meiosis-related genes in somatic cells reflects the difference in their DNA methylation status in which somatic cells derived from *Pcgf6*-null mice have been subjected to neither PRC1.6-mediated transcriptional repression nor Dnmt3B-mediated DNA methylation from one cell embryo. Aside from the possible role of PRC1.6 in somatic cells, it is clear that PRC1.6 has important roles as a safeguarding system that masks the potential of ESCs for their ectopic onset of meiosis. MGA functions as a scaffolding component in PRC1.6. Furthermore, MGA is involved in recruiting the whole complex to genomic DNA in a sequence-specific manner by utilizing its two distinct DNA binding domains. In the current study, we explored the roles of these two DNA-binding domains of Mga in transcriptional regulation in ESCs. Our data demonstrated that both DNA binding domains of Mga are functionally important by controlling rather distinct gene sets to each other, enabling them to regulate many genes in combination. Among the derepressed genes in ΔbHLHZ ESCs, genes that bear *de novo* genomic binding sites for Pcgf6 showed much more pronounced transcriptional activation than genes that do not. Given that genomic binding of Pcgf6 faithfully represents the binding of PRC1.6 in the genome, our ChIP-seq data for histone modifications indicated that the former gene set is epigenetically more activated because of the impairment of the function of PRC1.6 by the deletion of the bHLHZ domain of Mga. However, there was no significant difference in the magnitude of transcriptional activation because of the presence of *de novo* Pcgf6-binding sites among the activated genes in ΔT ESCs. ChIP-seq analyses for histone modifications also showed no (H3K27me3) or only subtle (H2AK119ub and H3K27Ac) differences in the epigenetic states of these genes because of the presence of Pcgf6-binding sites. A *de novo* sequence motif search around publicly reported Pcgf6-binding sites of specifically activated genes in ΔT ESCs revealed that, while the T-box sequence is barely identified, the E2F6 motif is identified as a much more prominent motif. Therefore, we assume that statistically no or only modest alterations in histone modifications among genes activated in ΔT ESCs, even in the presence of Pcgf6, may be attributed to the substantial contribution of activation that is secondary in nature, e.g., T-box deletion did not lead to complete disruption of PRC1.6, but caused a conformational change in the complex such that functions of E2F6 and/or certain other components of the complex were moderately affected. This notion is in accordance with a previous demonstration by Stielow et al.(2018) showing that E-box and E2F6 binding motif, but not T-box, were identified as prevailing motifs for the association with PRC1.6 in the genome, while the T-box sequence was identified only on the E2F6-null background.

Our data clearly demonstrated that deletion of the bHLHZ domain that renders Mga non-interactive with Max boosts the expression of a fairly similar set of meiosis-related genes activated in *Max*-KO ESCs. However, similarities in the expression profiles of the two cell-specific genes between *Max*-KO ESCs and ΔbHLHZ ESCs were rather limited in which two cell-specific genes was generally elevated in *Max*-KO ESCs irrespective of their relation to meiosis, whereas only a subset of those bearing meiosis-related characteristics were restrictively activated in ΔbHLHZ ESCs. Therefore, at least profound activation of two cell-specific genes that are unrelated to meiosis in *Max*-KO ESCs such as *Zscan4* did not reflect the impairment of the function of PRC1.6, but reflected the impairment of some other functions of Max. Interestingly, our data revealed that some marginal similarities in the activation profile of genes with neither meiosis-related characteristics nor *de novo* Pcgf6-binding sites were evident between *Max*-KO and ΔT ESCs. Furthermore, our qPCR analyses revealed that Dux, which is regarded as a master regulator of two cell-specific genes and its representative downstream genes, i.e., *Zscan4* (Cossec et al., 2018; De Iaco et al., 2017; Hendrickson et al., 2017), was activated in ΔT ESCs to some extent. This indicated that certain molecular mechanisms that repress the expression of two cell-specific genes were inactivated in ΔT ESCs, but were not sufficient for bringing wide-range activation of two cell-specific genes like *Max*-KO ESCs. Since MYC has been recently shown to be crucially involved in regulating transcription from two cell-specific genes(Fu et al., 2019), it is tempting to speculate that differences in the magnitude of activation of two cell-specific genes between *Max*-KO and ΔT ESCs are attributed to the differences in MYC activity, which is inactive and active in the former and the latter ESC lines, respectively.

*Stra8* is an essential gene in meiotic onset in germ cells (Anderson et al., 2008; Baltus et al., 2006; Dokshin et al., 2013; Mark et al., 2008) and our previous study demonstrated that it is also essential for ectopic meiosis in ESCs (Suzuki et al., 2016). Although the molecular basis of Stra8 that promotes meiosis has remained largely obscure until recently, the identification of Meiosin as indispensable partner for Stra8 drastically changed this elusive situation (Ishiguro et al., 2020). Since the molecular basis of positive and negative regulation of meiosis sustained by RA–Stra8 signaling and PRC1.6, respectively, are independently fairly well understood, there a base from which to explore the molecular mechanisms that regulate these two opposing regulations in an inversely correlated manner so that one dominates over the other depending on the cell status. It is known that PRC1.6 does not bind to *Stra8* and therefore does not influence its expression directly. However, our data clearly demonstrate that *Meiosin* is a target of PRC1.6 whose repressed state is influenced by the integrity of Mga protein, in particular, by that of the bHLHZ domain. Therefore, in conjunction with a previous report (Ishiguro et al., 2020), our data indicate that Meiosin serves as a critical molecule in the transition from mitosis to meiosis in which PRC1.6 renders the RA–Stra8 signaling axis inactive via the repression of *Meiosin* in the mitotic phase. The relative dominance of the two opposing regulations are reversed upon meiotic onset.

Although we are far from gaining a complete understanding of the molecular mechanisms that block the potential for ectopic meiotic onset of ESCs, we hope that the data presented in this study will serve as a positive step towards achieving this goal. We also anticipate that our data will help reveal the molecular mechanisms that inactivate the function of PRC1.6 at the onset of meiosis in germ cells.

## ACKNOWLEDGMENTS

We are indebted to Kazumi Ubukata for his technical assistance. This work was supported in part by the Ministry of Education, Culture, Sports, Science and Technology (MEXT), Japan. K.U. and A.O. are recipients of grants from the Japan Society for the Promotion of Science (JSPS) KAKENHI with grant numbers of 20K16147 and 19H03426, respectively. This work was also supported in part by the Research Fellowship for Young Scientists (DC2) from JSPS to Y.K. We thank H. Nikki March, PhD, from Edanz Group (https://en-author-services.edanzgroup.com/) for editing a draft of this manuscript.

## Notes

### Competing Interest Statement

The authors have declared no competing interest.

