## Supplemental Fig. 1-5 and Tables 1-4 for "Two DNA binding domains of Mga act in combination to suppress ectopic activation of meiosis-related genes in mouse embryonic stem cells"

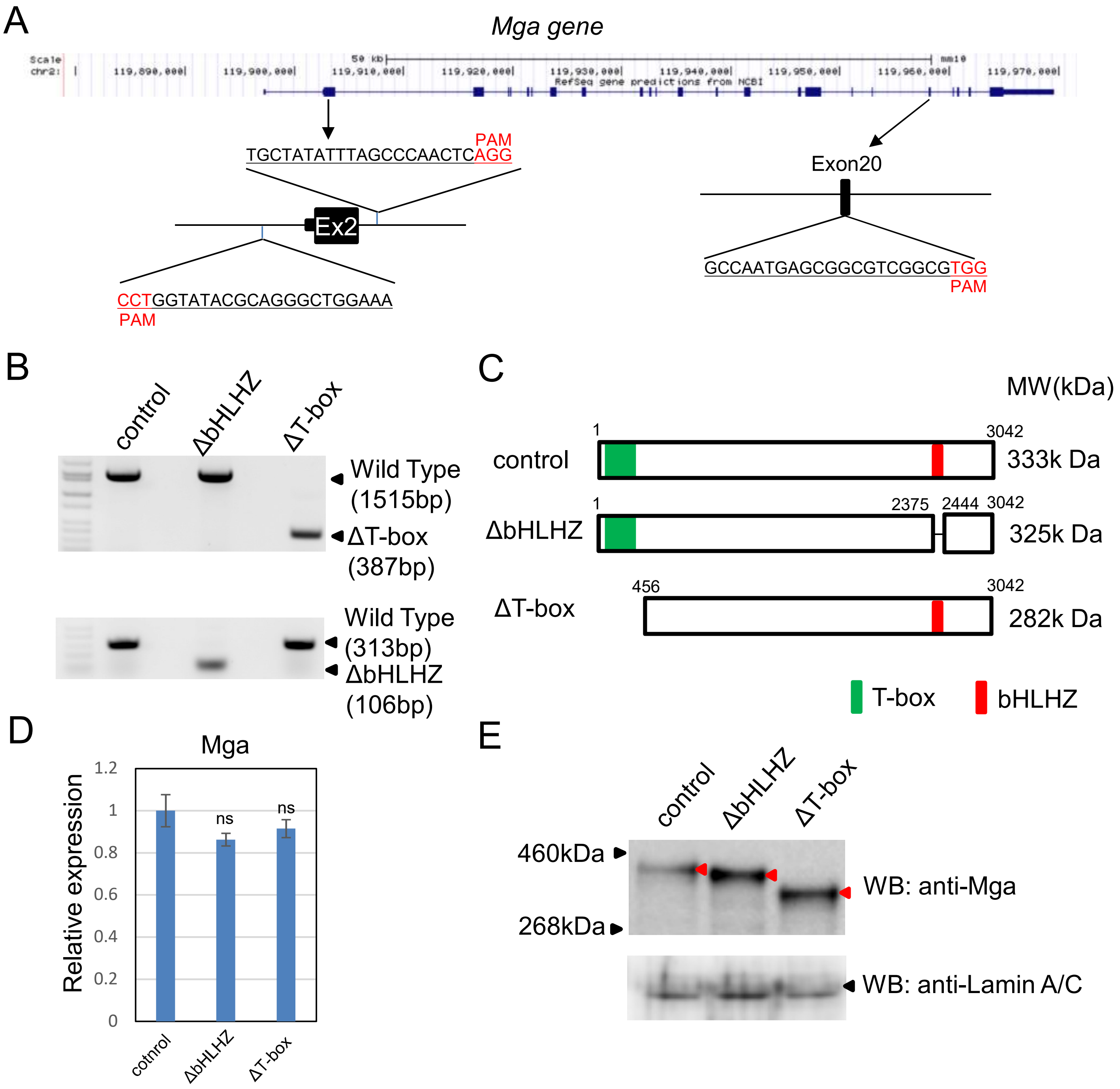


**Figure S1. Generation of two distinct mutant ESCs that produce Mga lacking either T-box or bHLHZ domain.** (A) Schematic representation of the strategy for generating two *Mga*-mutant ESCs lacking either T-box-encoding exon 2 or bHLHZ domain-encoding exon 20. These exons were deleted in ESCs by means of the CRISPR-Cas9 system using the indicated oligonucleotides. (B) RT-PCR for screening of candidate ESC clones in which exon 2 (left panel) or exon 20 was successfully removed. (C) Schematic diagrams of wild-type and two mutants of Mga protein. Green and red boxes indicate T-box and bHLHZ domains, respectively. (D) Quantification of *Mga* mRNA in parental and *Mga*-mutant ESC lines by qPCR. Expression levels of *Mga* mRNA in parental ESCs were arbitrarily set to one. Data represents mean ± standard deviation of three independent experiments. Student's t-test was conducted to examine statistical significance. ns, not significant. (E) Western blot analyses of Mga in parental and distinct *Mga*-mutant ESC lines. Black and red arrows indicate specific protein bands for LaminA/C and wild-type Mga or its derivatives, respectively.

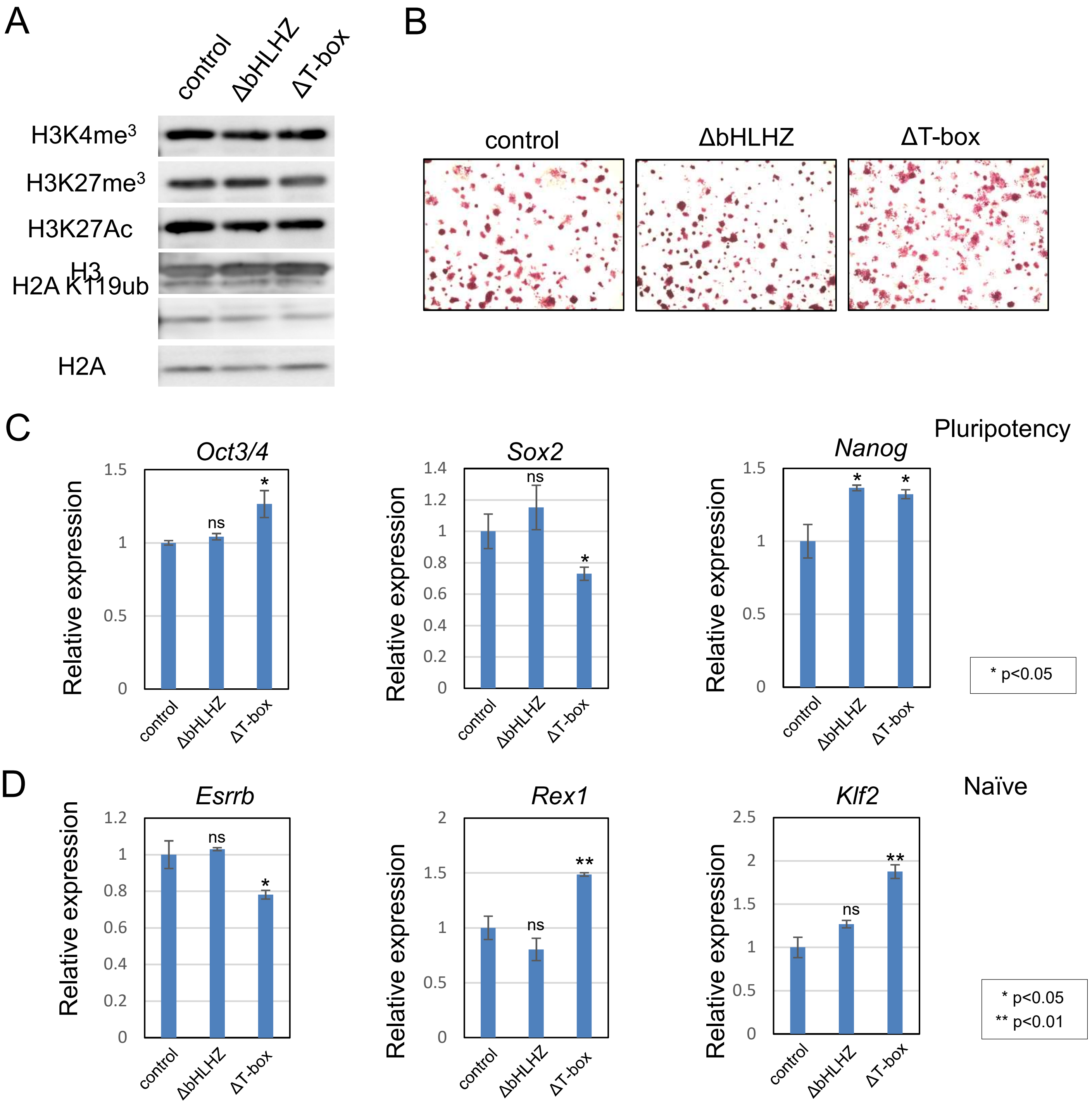


**Figure S2. Examination of global histone modification levels and indicators of the undifferentiated state in parental and *Mga*-mutant ESCs.** (A) Western blot analyses of histone modifications. Antibodies against histone H2A and H3 and their derivatives with the indicated post-translational modifications were used to determine their relative amounts among parental and *Mga*-mutant ESC lines. (B) Alkaline phosphatase staining of parental and *Mga*-mutant ESC lines. (C) Examination of the expression levels of pluripotency marker genes. Expression levels of *Oct4*, *Sox2*, and *Nanog* in parental and *Mga*-mutant ESCs were quantified as representative pluripotency marker genes by qPCR. Data represents mean ± standard deviation of three independent experiments. Student's t-test was conducted to examine statistical significance. ns, not significant. (D) Examination of expression levels of naïve marker genes. Expression levels of *Esrrb*, *Rex1*, and *Klf2* genes were quantified as in C as representative naïve marker genes. Data were treated statistically as in C.

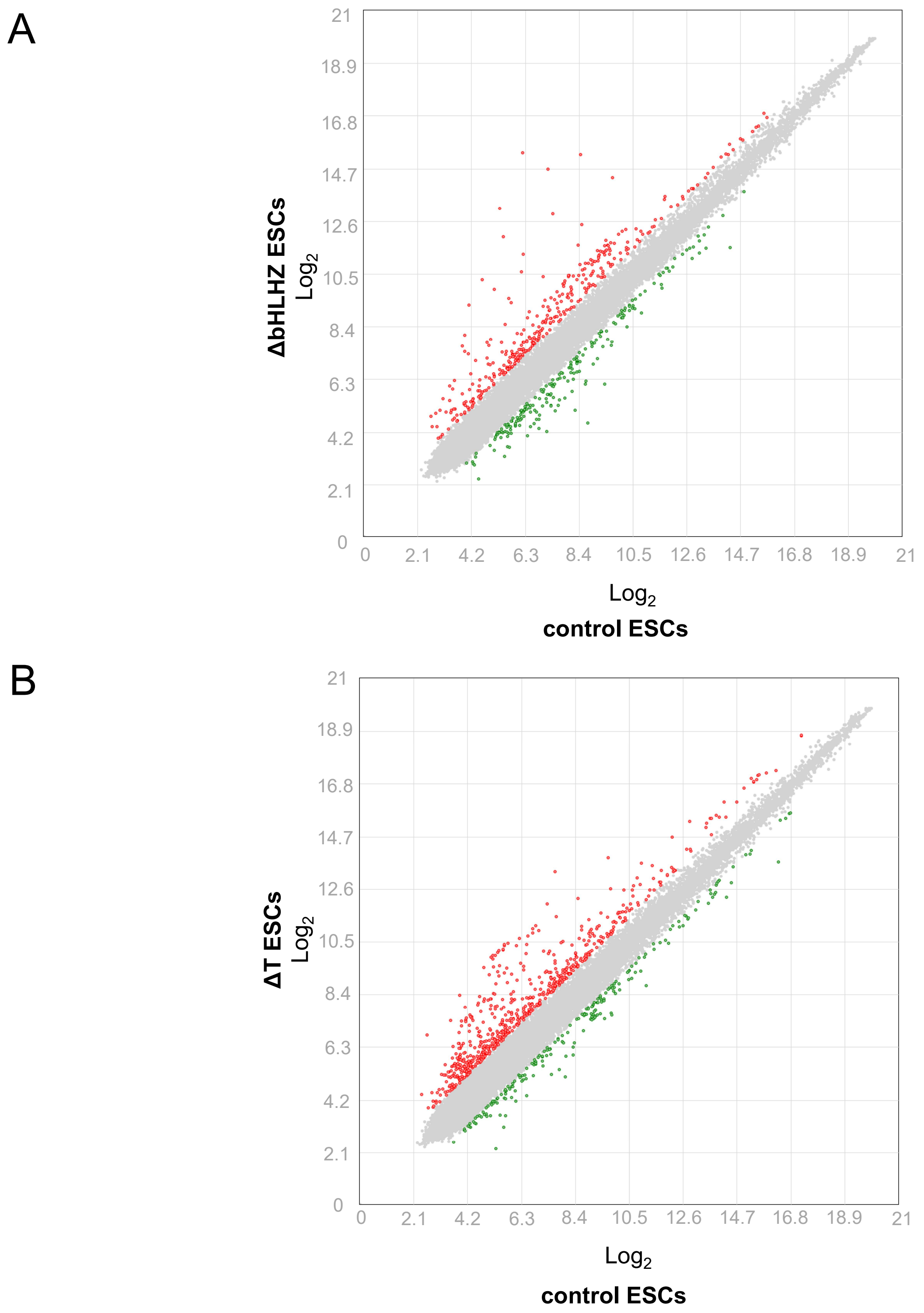


**Figure S3. Examination of the effects of *Mga* mutations on global expression profiles in ESCs.** (A) Scatter plots of DNA microarray data from parental and ΔbHLHZ ESCs. Genes showing more than 2-fold higher expression values in ΔbHLHZ ESCs and parental ESCs are highlighted by red and green dots, respectively. (B) Scatter plots of DNA microarray data from parental and ΔT ESCs. Genes showing more than 2-fold higher expression values in ΔT ESCs and parental ESCs are highlighted by red and green dots, respectively.

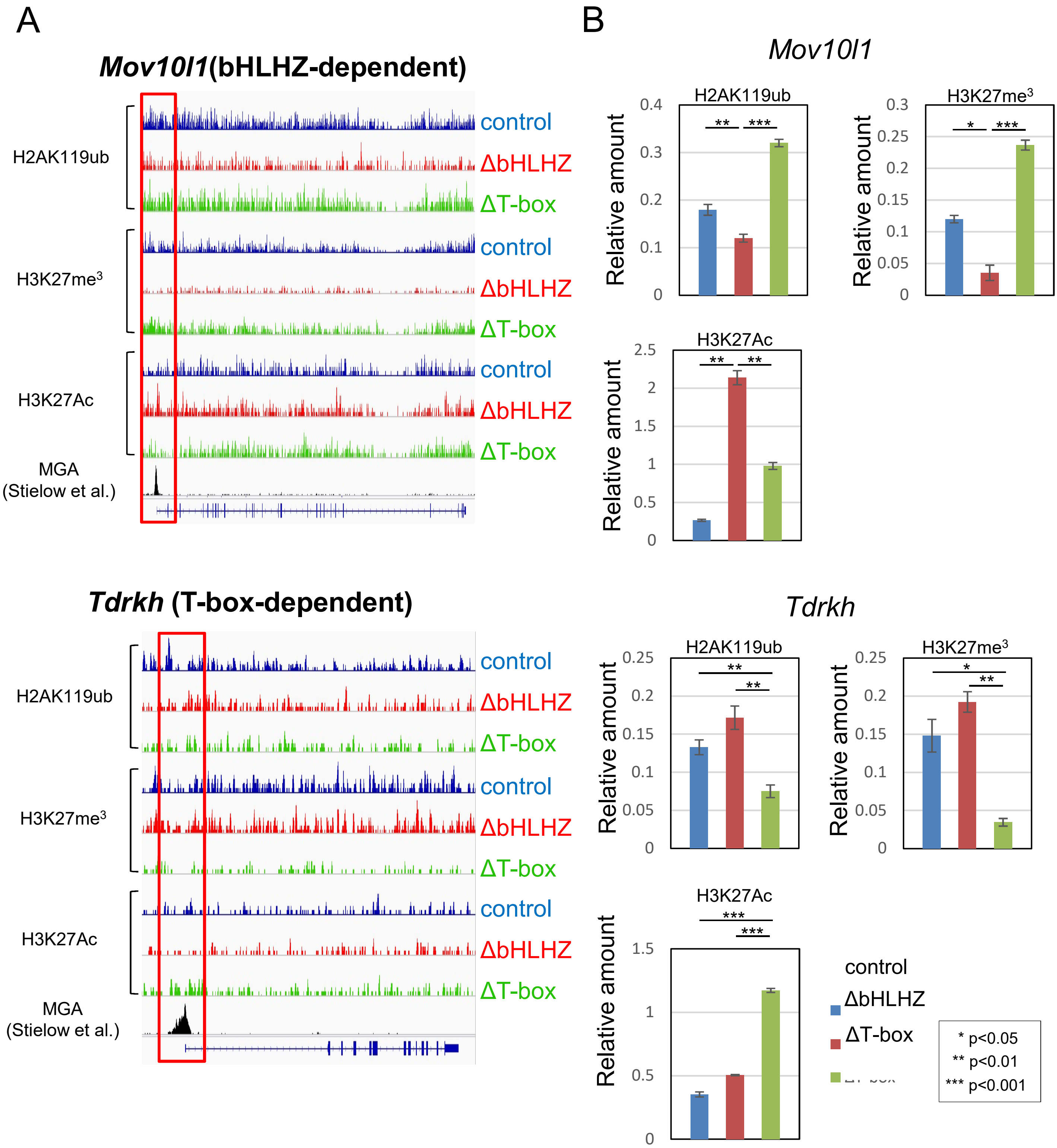


**Figure S4. Examination of histone modification levels of *Mov10l1* and *Tdrkh* genes in ΔbHLHZ and ΔT ESCs. (A):** ChIP-seq profiles of histone modifications. *Mov10l1* and *Tdrkh* were selected as representative genes subjected to bHLHZ- and T-box-dependent repression in ESCs. Mga binding profiles in these genes were constructed using publicly reported ChIP-seq data (Stielow et al., 2018). (B) Examination of histone modifications by locus-specific ChIP-qPCR. ChIP-qPCR analyses of histone modifications of H2AK119ub, H3K27me$^3$, and H3K27Ac were performed at publicly reported Mga-binding sites of *Mov10l1* and *Tdrkh*. Data represents mean ± standard deviation of three independent experiments. Student's t-test was conducted to examine statistical significance.

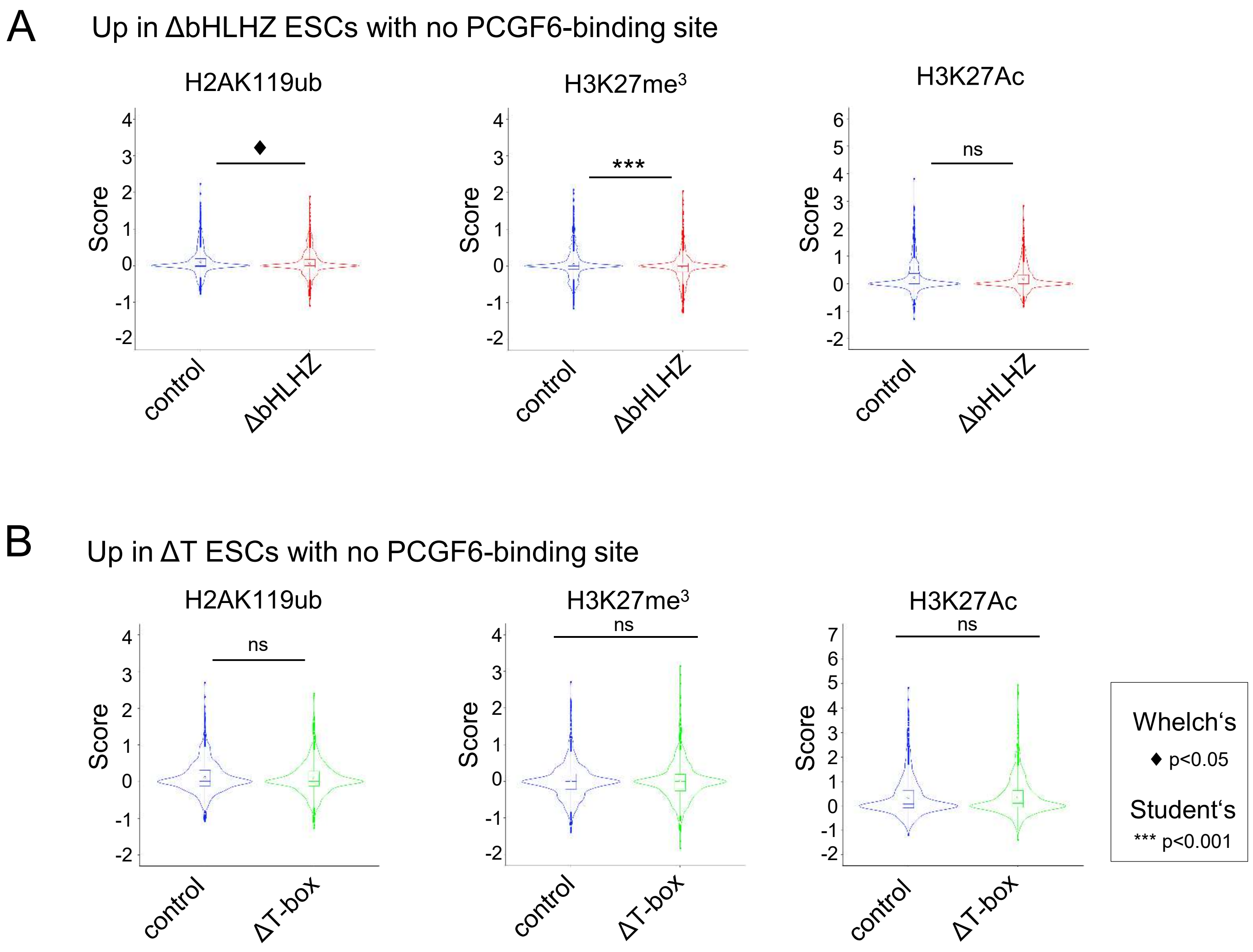


**Figure S5. Examination of epigenetic alterations in the activated genes with no *de novo* genomic binding sites for Pcgf6 in *Mga*-mutant ESCs.** (A) Histone modification analyses of the activated genes in ΔbHLHZ ESCs with no publicly reported Pcgf6-binding sites. ChIP-seq data were used to examine the levels of alterations in histone modifications (H2AK119ub, H3K27me$^3$, and H3K27Ac) of genes that were activated in ΔbHLHZ ESCs, but that do not bear publicly reported Pcgf6-binding sites. Genomic regions around a transcription start site of each gene (5 kb) were used to determine the score for histone modification levels. Statistical analyses were conducted as in Figure 2D. (B) Histone modification analyses of the activated genes in ΔT ESCs with no publicly reported Pcgf6-binding sites. The scores for histone modifications were obtained as in A with genes activated in ΔT ESCs with no *de novo* Pcgf6-binding sites. Data were treated statistically as in Figure 2D.

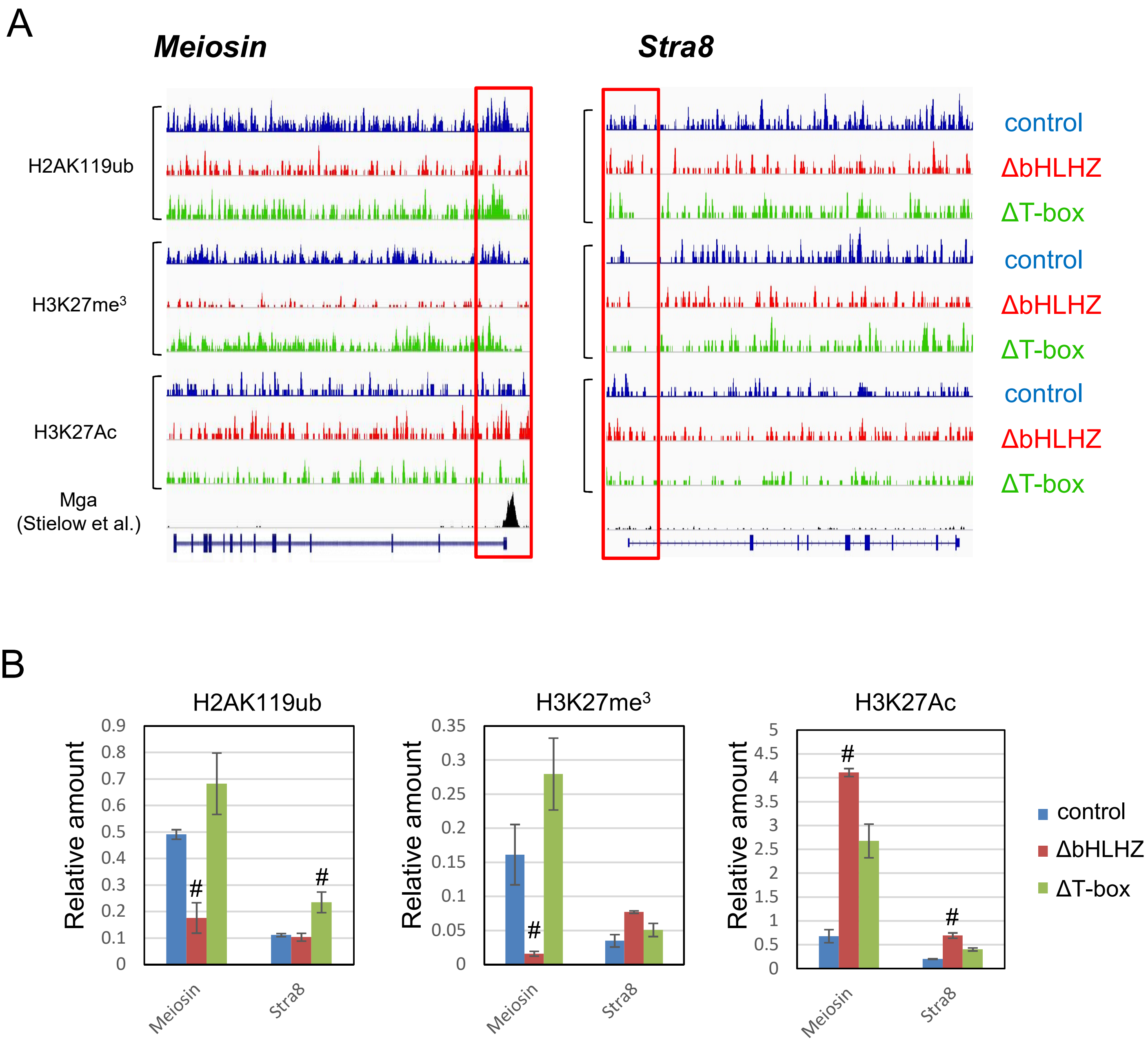


**Figure S6. Examination of the histone modification levels of *Meiosin* and *Stra8* genes in ΔbHLHZ and ΔT ESCs.** (A) ChIP-seq profiles of histone modifications around *Meiosin* and *Stra8* genes. Publicly reported ChIP-seq data (Stielow et al., 2018) were used for constructing a Mga binding profile for *Meiosin*, while there was no evidence of binding of Mga on *Stra8*. (B) Examination of histone modifications by locus-specific ChIP-qPCR. ChIP-qPCR analyses of histone modifications (H2AK119ub, H3K27me$^3$, and H3K27Ac) were performed for *Meiosin* and *Stra8* at publicly reported Mga binding sites and transcription start sites, respectively. Data obtained with DNA recovered from samples that were not subjected to immunoprecipitation were arbitrarily set to one. Data represents mean ± standard deviation of three independent experiments. Student's t-test was conducted to examine statistical significance.

### Table S1. Sequences of oligonucleotides

#### Oligonucleotides for genetic manipulation of *Mga* loci by CRISPR/Cas9

| Forward | Reverse |
|---|---|
| ***Mga* intron 1 (forΔT-box)** | |
| 5'-caccgTTTCCAGCCCTGCGTATACC-3' | 5'-aaacGGTATACGCAGGGCTGGAAAc-3' |
| ***Mga* intron 2 (forΔT-box)** | |
| 5'-caccgTGCTATATTTAFCCCAACTC-3' | 5'-aaacGAGTTGGGCTAAATATAGCAc-3' |
| ***Mga* Exon20 (for ΔbHLH)** | |
| 5'-caccGCCAATGAGCGGCGTCGGCG-3' | 5'-aaacCGCCGACGCCGCTCATTGGC-3' |

#### Genotyping

| Forward | Reverse |
|---|---|
| ***Mga* Exon 1 – Exon 3** | |
| 5'-CAGCTGAAGCGGACACACG-3' | 5'-CTGTTATCCCTTCTGGGCATTCACC-3' |
| ***Mga* Exon 19 - Exon 21** | |
| 5'-GTGTCATAGCCATATCTCTGCAGATG-3' | 5'-CTGCCCTATCAATTTATCTGCCTGG-3' |

#### Quantitative PCR

| Forward | Reverse |
|---|---|
| ***Mga*** | |
| 5'- TGGTAGCGGTTTCTCCTGCTC-3' | 5'- GTCAAGTGGCTTGCCATCAGG-3' |
| ***Mov10l1*** | |
| 5'- GAGAAGCCCATCGTGGTTCAGC-3' | 5'- TTTCCACCTGCTTCCGATAGGG -3' |
| ***Boll*** | |
| 5'- ACGGTGGCTGTGTTCCTCCT -3' | 5'- AATTGGCTCAGGCTGCATCATAGGT -3' |
| ***Tdrkh*** | |
| 5'- TTCTGGTGCCCAGAGCAGTC -3' | 5'- GGCTGCGGGAACCAATGATTTG -3' |
| ***Tex12*** | |
| 5'- GAGAAGGATTTGAGCGATATGAGCAAGG -3' | 5'- CTGTAAACCTCTGCTTCAGGAACTCTC -3' |
| ***Stag3*** | |
| 5'- GCAAGCAGCTGACCCGACT -3' | 5'- GGTCCAACCCATGCTCACTGG -3' |
| ***Fkbp6*** | |
| 5'- TGACAAAAGGAACGCCAAGGCC -3' | 5'- CTGCTTCCACAGGGAGCAAACA -3' |

| | |
|---|---|
| ***Meiosin*** | |
| 5’- CATTGACATGACCAAGGCCTTGC -3’ | 5’-TGGAGGGAGTGGAGTGTTGCT-3’ |
| ***Hormad2*** | |
| 5’-AGGCCACCAGAATGCTCGAT-3’ | 5’-TGTCACTGGTTCGCTGACCTT-3’ |

**ChIP-qPCR**

| Forward | Reverse |
|---|---|
| ***Mov10l1*** | |
| 5’- GCCACCCTCTCTTCCACACA-3’ | 5’- ACGAGAGGTTGGCGGTTAGC-3’ |
| ***Tdrkh*** | |
| 5’- GGAGGACCGAGGACTCCAGA-3’ | 5’- CCCACGTCCACAGCCAGTTA-3’ |
| ***Meiosin*** | |
| 5’- AGCAGAAGTCGAGGCTTAACGC-3’ | 5’- TGACTCTGACGACTCCTCACCAT-3’ |
| ***Stra8*** | |
| 5’- TGTGATTGGTTCGCAGCCTG-3’ | 5’- CTGCCCGTCGCAGAATAAGAAG-3’ |

### Table S2 Materials used in this study

##### Antibodies

| Antigen | Manufacturer | Catalog No. |
|---|---|---|
| histone H3 | Cell Signaling | 9715S |
| histone H3K27me$^3$ | Millipore | 07-499 |
| histone H3K27Ac | abcam | ab4729 |
| histone H2A | Cell Signaling | 12349S |
| histone H2AK119ub | Cell Signaling | 8240S |
| lamin A/C | SANTA CRUZ | sc-20681 |
| Stra8 | abcam | ab40602 |
| Sycp3 | abcam | ab97672 |

##### TaqMan probes used for qPCR

| Gene Symbol | Probe ID |
|---|---|
| *Stra8* | Mm00486473_m1* |
| *Sycp3* | Mm00488519_m1 |
| *Taf7l* | Mm00459354_m1 |
| *Slc25a31* | Mm00617754_m1 |
| *Dazl* | Mm03053726_s1 |
| *Dux* | Mm03053726_s1; |
| *Zscan4* | Mm01234988_g1 |
| *Oct4* | Mm00658129_gH |
| *Sox2* | Mm00488369_s1; |
| *Nanog* | Mm02019550_s1 |
| *Esrrb* | Mm00442411_m1 |
| *Rex1* | Mm03053975_g1; |
| *Klf2* | Mm01244979_g1 |
| *Gapdh* | Mm99999915_g1 |

*All of TaqMan probes are from Thermo Fisher Scientific

### Table S3

#### List of Up-regulated gnes in ΔbHLHZ and/or ΔT ESCs with information of *de novo* PCGF6-binding site

Fold change

>50

0.1>

| | | | Δ bHLHZ | Δ T |
|---|---|---|---|---|
| ΔHLHZ ≧ 2 ; ΔT < 2 | PCGF6-binding site ( + ) | Mov10l1 | 740.49 | 1.93 |
| | | Hormad1 | 223.31 | 0.84 |
| | | Dazl | 45.76 | 1.05 |
| | | Sycp3 | 30.18 | 0.98 |
| | | Tdrd9 | 16.55 | 1.01 |
| | | Boll | 13.79 | 0.66 |
| | | Rnf17 | 12.16 | 1.64 |
| | | Mei1 | 12.07 | 0.94 |
| | | Nxf2 | 9.68 | 1.26 |
| | | Tdrd1 | 8.50 | 1.01 |
| | | Adad2 | 7.50 | 1.06 |
| | | Casp7 | 5.33 | 0.68 |
| | | Micu2 | 5.33 | 1.18 |
| | | Tcam1 | 5.17 | 0.92 |
| | | Gm10406 | 4.81 | 0.97 |
| | | 2610042L04Rik | 4.54 | 1.04 |
| | | Gm3411 | 4.40 | 1.01 |
| | | Ddx25 | 3.96 | 1.30 |
| | | Mga | 3.75 | 1.03 |
| | | Akap11 | 3.72 | 0.64 |
| | | Elovl4 | 3.65 | 0.63 |
| | | Ptprs | 3.61 | 1.19 |
| | | Cryl1 | 3.60 | 0.83 |
| | | Ripk3 | 3.49 | 1.24 |
| | | Pus7l | 3.28 | 1.00 |
| | | Tmem185b | 3.28 | 1.24 |
| | | Cyct | 3.19 | 1.33 |
| | | Ciart | 3.17 | 1.21 |
| | | Dgat2 | 3.16 | 1.60 |
| | | Spata22; Olfr20 | 2.87 | 1.15 |
| | | 4930447C04Rik | 2.86 | 1.26 |
| | | Fam208b | 2.79 | 0.52 |
| | | Paqr9 | 2.79 | 0.99 |
| | | Dmrt1 | 2.74 | 1.56 |
| | | Tsnax | 2.62 | 0.76 |
| | | Eif5a2 | 2.60 | 0.70 |
| | | Nefh | 2.56 | 1.06 |
| | | Prkar2b | 2.56 | 1.59 |
| | | Smc1b | 2.54 | 1.16 |
| | | Ppp1r3d | 2.50 | 1.10 |
| | | Pmaip1 | 2.49 | 1.35 |
| | | Pom121 | 2.45 | 1.95 |
| | | Gtf2f2 | 2.44 | 0.84 |
| | | Suclg2 | 2.43 | 1.84 |
| | | Trim44 | 2.35 | 1.21 |
| | | Gm3727 | 2.33 | 0.95 |

| | | | | |
|---|---|---|---|---|
| | | Fam183b | 2.31 | 1.29 |
| | | Chmp7 | 2.28 | 0.65 |
| | | Ipo4 | 2.28 | 0.66 |
| | | Igf2r | 2.25 | 1.36 |
| | | Gm128 | 2.24 | 1.16 |
| | | Gm3115 | 2.23 | 0.89 |
| | | Fam217b | 2.21 | 1.23 |
| | | Cpeb1 | 2.18 | 1.30 |
| | | Uqcc2 | 2.18 | 1.20 |
| | | Zfp951 | 2.13 | 1.53 |
| | | Mfsd1 | 2.12 | 1.00 |
| | | Ngdn | 2.05 | 0.60 |
| | PCGF6-binding site (-) | Hpse2 | 16.66 | 1.08 |
| | | Gm3298 | 5.60 | 1.23 |
| | | Gm3468 | 5.46 | 0.93 |
| | | Gm3383; Gm3194 | 5.39 | 0.88 |
| | | Gm2897 | 5.14 | 0.93 |
| | | Gm5796; Gm3095; Gm8108 | 4.95 | 0.95 |
| | | Gm17651 | 4.92 | 0.58 |
| | | Gm3383; Gm10021 | 4.91 | 1.00 |
| | | Gm3591 | 4.91 | 0.95 |
| | | Gm3667 | 4.90 | 1.09 |
| | | Gm3264; Gm3173 | 4.85 | 1.11 |
| | | Gm2956 | 4.67 | 1.01 |
| | | Gm3020; Gm10409 | 4.65 | 0.89 |
| | | Gm3317; Gm3488 | 4.64 | 0.94 |
| | | Gm2897; Gm3005 | 4.63 | 0.95 |
| | | Gm3500 | 4.62 | 0.94 |
| | | Gm10128 | 4.59 | 1.11 |
| | | Gm3558 | 4.58 | 0.89 |
| | | Gm2974 | 4.50 | 0.94 |
| | | Gm10340 | 4.48 | 0.99 |
| | | Gm3239 | 4.46 | 1.02 |
| | | Gm3636; Gm3264 | 4.44 | 1.13 |
| | | Gm2237 | 4.40 | 0.96 |
| | | Gm10021 | 4.35 | 1.12 |
| | | Gm3264 | 4.33 | 1.09 |
| | | Gm3002 | 4.31 | 0.86 |
| | | Gm3500; Gm3373 | 4.24 | 0.78 |
| | | Mir669b | 4.20 | 0.43 |
| | | Gm8237 | 4.11 | 1.00 |
| | | Gm3696 | 3.83 | 0.87 |
| | | Gm8281 | 3.76 | 0.69 |
| | | Gm8206 | 3.75 | 1.00 |
| | | Gm3739 | 3.74 | 0.91 |
| | | Neto2 | 3.66 | 1.24 |
| | | Hdac6 | 3.08 | 1.64 |
| | | Hbp1 | 3.01 | 0.97 |
| | | Gm3248 | 2.97 | 0.81 |
| | | Gm8279 | 2.94 | 0.82 |
| | | Gm3029 | 2.92 | 0.90 |
| | | Gm7972 | 2.89 | 0.98 |
| | | Gm8159 | 2.89 | 1.01 |

| | | | | |
|---|---|---|---|---|
| | | Gm10338 | 2.87 | 0.74 |
| | | Gm8297 | 2.86 | 0.95 |
| | | Gm5795; Gm10413 | 2.85 | 0.79 |
| | | Gm3187 | 2.83 | 0.82 |
| | | Efna3 | 2.82 | 1.20 |
| | | Gm3453 | 2.82 | 0.84 |
| | | Gm5795 | 2.82 | 0.75 |
| | | Gm3242 | 2.78 | 0.99 |
| | | Gm3642; LOC100504218 | 2.77 | 0.74 |
| | | Gm5797 | 2.76 | 0.69 |
| | | Irf2bp2 | 2.75 | 0.74 |
| | | Gm3512 | 2.73 | 0.82 |
| | | Gm3685 | 2.73 | 0.78 |
| | | Gm16440 | 2.72 | 1.35 |
| | | Gm2888 | 2.70 | 0.82 |
| | | Gm5795; Gm10413; Gm3012 | 2.69 | 0.74 |
| | | Extl3 | 2.68 | 0.84 |
| | | Gm21560 | 2.64 | 0.78 |
| | | Gm3476 | 2.60 | 0.75 |
| | | Klf4 | 2.58 | 0.83 |
| | | Mlc1 | 2.57 | 0.80 |
| | | Slc7a7 | 2.56 | 1.12 |
| | | Gm8362 | 2.51 | 0.98 |
| | | Gm5514 | 2.49 | 0.61 |
| | | Gm3182 | 2.45 | 0.66 |
| | | Gm3594 | 2.45 | 0.80 |
| | | Mphosph8 | 2.44 | 0.55 |
| | | Trim13 | 2.42 | 1.42 |
| | | Cdkn2d | 2.40 | 1.25 |
| | | Gm3138 | 2.40 | 0.84 |
| | | Gm6337 | 2.40 | 0.99 |
| | | Mir669h | 2.39 | 0.97 |
| | | Gm3149 | 2.38 | 0.88 |
| | | Gm3526 | 2.38 | 0.71 |
| | | Rnu6 | 2.38 | 0.90 |
| | | Tfcp2l1 | 2.36 | 0.99 |
| | | Gm3542 | 2.35 | 0.77 |
| | | Gm9603 | 2.35 | 0.77 |
| | | D830030K20Rik | 2.34 | 0.99 |
| | | Gm3182; Gm7876 | 2.34 | 0.70 |
| | | Trim36 | 2.34 | 1.00 |
| | | Gm8356 | 2.33 | 0.82 |
| | | Pim2 | 2.33 | 1.45 |
| | | Fgf5 | 2.30 | 0.85 |
| | | Gm8246 | 2.30 | 0.63 |
| | | Lss | 2.30 | 0.76 |
| | | Mtmr9 | 2.29 | 0.59 |
| | | Ipw; Snord116l1; Snord116l2; Snord116 | 2.28 | 1.71 |
| | | Ccar2 | 2.27 | 0.66 |
| | | Sorl1 | 2.27 | 1.01 |
| | | Gm3127 | 2.26 | 0.70 |
| | | Gm3460 | 2.25 | 0.68 |

| | | | | |
|---|---|---|---|---|
| | | Crip2 | 2.23 | 1.01 |
| | | Gm3339 | 2.23 | 0.71 |
| | | Gm8265 | 2.23 | 0.71 |
| | | Mtmr6 | 2.23 | 0.88 |
| | | Gm21103 | 2.20 | 0.88 |
| | | Als2cr11 | 2.19 | 0.88 |
| | | Jph1 | 2.18 | 1.06 |
| | | Sox15 | 2.17 | 0.75 |
| | | Cpsf4l | 2.16 | 1.37 |
| | | Gm3594; Gm3269 | 2.16 | 0.76 |
| | | Ints6 | 2.16 | 0.62 |
| | | Nlrp4a | 2.15 | 0.86 |
| | | Gm5458 | 2.13 | 0.78 |
| | | Zmym2 | 2.12 | 0.54 |
| | | Slc29a1 | 2.11 | 0.86 |
| | | Jag2 | 2.09 | 1.62 |
| | | Gm715 | 2.08 | 1.13 |
| | | Slc39a14 | 2.08 | 0.65 |
| | | Srgap2 | 2.08 | 1.57 |
| | | Gm16434 | 2.07 | 0.82 |
| | | Gm3424 | 2.07 | 0.72 |
| | | Gm6676 | 2.07 | 0.82 |
| | | Gm7831 | 2.07 | 1.17 |
| | | Slc35f5 | 2.06 | 1.16 |
| | | Fdft1 | 2.04 | 0.63 |
| | | Gm13422 | 2.04 | 1.41 |
| | | Fabp5 | 2.03 | 0.88 |
| | | Gm11343 | 2.03 | 1.12 |
| | | Gm3099 | 2.03 | 0.95 |
| | | Sema6a | 2.03 | 0.73 |
| | | Gm3159 | 2.02 | 0.73 |
| | | Ints9 | 2.02 | 0.57 |
| | | Nphp3 | 2.02 | 0.95 |
| | | Ccdc25 | 2.01 | 0.78 |
| | | Rdh11 | 2.01 | 0.72 |
| | | Supt16 | 2.01 | 0.48 |
| | | Esco2 | 2.00 | 0.67 |
| | | Prkcg | 2.00 | 1.00 |
| ΔHLHZ and ΔT ≧ 2 | PCGF6-binding site （+） | Gm4969 | 196.47 | 2.27 |
| | | Ddx4 | 127.53 | 2.69 |
| | | Syce1 | 50.33 | 3.81 |
| | | Ctcfl | 49.66 | 2.83 |
| | | Fkbp6 | 39.37 | 13.93 |
| | | Sycp2 | 29.56 | 3.76 |
| | | Sycp1 | 26.25 | 2.10 |
| | | Stag3 | 17.82 | 13.36 |
| | | Piwil2 | 15.93 | 2.68 |
| | | Mael | 10.89 | 2.69 |
| | | Gpx6 | 8.32 | 4.99 |
| | | Taf7l | 5.66 | 2.10 |
| | | Prss42 | 3.89 | 6.32 |
| | | Tdrd12 | 3.85 | 3.10 |
| | | Prr19 | 3.74 | 2.03 |

| | | | | |
|---|---|---|---|---|
| | | Slc25a31 | 3.40 | 2.30 |
| | | Fam178b | 3.18 | 2.16 |
| | | Stk31 | 2.97 | 2.10 |
| | | Sec1 | 2.93 | 2.43 |
| | | Rad50 | 2.44 | 2.06 |
| | | Ddah1 | 2.36 | 3.03 |
| | | Tex11 | 2.16 | 3.76 |
| | | Fam96a | 2.12 | 2.06 |
| | | Hormad2 | 2.10 | 17.75 |
| | PCGF6-binding site（-） | Snord116l1; Snord116l2; Snord116 | 5.63 | 2.41 |
| | | Stra8 | 5.39 | 2.57 |
| | | Tex15 | 4.12 | 2.17 |
| | | P2rx7 | 3.69 | 21.56 |
| | | Ece1 | 3.20 | 2.14 |
| | | Ahsg | 2.98 | 2.27 |
| | | Gm17482 | 2.78 | 2.50 |
| | | Lgals1 | 2.72 | 4.76 |
| | | Gm13503 | 2.42 | 2.14 |
| | | Zbtb32 | 2.39 | 3.12 |
| | | Ephx2 | 2.26 | 2.13 |
| | | Trp53inp1 | 2.25 | 2.31 |
| ΔT ≧ 2; ΔHLHZ ＜ 2 | PCGF6-binding site（+） | Zcwpw1 | 0.60 | 52.53 |
| | | Tdrkh | 0.84 | 22.33 |
| | | Tex12 | 1.15 | 14.33 |
| | | Tex19.1 | 1.53 | 7.91 |
| | | Gpat2 | 1.17 | 6.63 |
| | | Bin2 | 0.91 | 6.48 |
| | | Hist1h2ba | 0.65 | 6.35 |
| | | Hist1h2aa | 0.36 | 5.46 |
| | | Rpl10l | 0.63 | 4.82 |
| | | Peg10 | 1.15 | 4.47 |
| | | D1Pas1 | 0.49 | 4.34 |
| | | Rsrp1 | 0.71 | 3.99 |
| | | Liph | 0.39 | 3.59 |
| | | Plk2 | 1.89 | 3.39 |
| | | Wbp2nl | 0.91 | 3.21 |
| | | Syce3 | 1.54 | 3.13 |
| | | Csrnp2 | 1.37 | 3.12 |
| | | Gadd45b | 0.71 | 3.04 |
| | | Cenpm | 0.85 | 3.00 |
| | | Ndrg1 | 0.59 | 2.79 |
| | | Zfp677 | 0.72 | 2.66 |
| | | Plekhb2 | 0.59 | 2.64 |
| | | Slc47a1 | 0.93 | 2.63 |
| | | Aard | 1.04 | 2.54 |
| | | Klhl22 | 0.76 | 2.52 |
| | | Colec12 | 0.87 | 2.49 |
| | | Commd5 | 0.91 | 2.41 |
| | | Lhpp | 1.34 | 2.35 |
| | | Mlf1 | 0.91 | 2.29 |
| | | Psap | 0.47 | 2.29 |
| | | Tmem181a | 1.77 | 2.28 |
| | | Carhsp1 | 1.49 | 2.26 |

| | | | | |
|---|---|---|---|---|
| | | Gtsf1 | 0.79 | 2.26 |
| | | Pla2g6 | 1.14 | 2.24 |
| | | Calcoco1 | 1.21 | 2.21 |
| | | Sord | 1.61 | 2.21 |
| | | Xpnpep3 | 0.73 | 2.21 |
| | | Apobec2 | 0.62 | 2.20 |
| | | Prss50 | 1.53 | 2.20 |
| | | Ccno | 1.16 | 2.19 |
| | | Tuba3a | 0.52 | 2.19 |
| | | Rec114 | 1.22 | 2.17 |
| | | Rnf213 | 0.73 | 2.17 |
| | | Mtss1 | 0.80 | 2.10 |
| | | Ldah | 1.40 | 2.09 |
| | | Acaa2 | 1.37 | 2.08 |
| | PCGF6-binding site (-) | Gm364 | 1.39 | 28.06 |
| | | Zscan4f | 0.22 | 26.22 |
| | | Obox4-ps11 | 0.59 | 24.20 |
| | | Zscan4e | 0.27 | 19.86 |
| | | Zscan4b | 0.19 | 19.71 |
| | | Rhox5 | 0.24 | 18.60 |
| | | LOC215866 | 1.13 | 18.37 |
| | | Zscan4a | 0.23 | 17.26 |
| | | Zscan4c | 0.30 | 15.96 |
| | | Zscan4d | 0.31 | 13.53 |
| | | Obox4-ps8 | 0.61 | 13.32 |
| | | Obox4-ps36 | 0.65 | 12.64 |
| | | Obox4-ps6 | 0.99 | 11.62 |
| | | Gm4340; Gm21312; Gm20765; Gm21293; Gm6763; Gm8764; Gm21304 | 0.51 | 10.59 |
| | | Gm6468 | 1.08 | 10.55 |
| | | Gm7647 | 0.61 | 10.02 |
| | | Gm6502 | 0.94 | 9.79 |
| | | Gm6523 | 0.83 | 9.59 |
| | | Rhox13 | 1.14 | 9.58 |
| | | Gm2016 | 0.51 | 9.24 |
| | | Gm6348 | 0.88 | 9.17 |
| | | Gm21312; Gm20765; Gm21293; Gm6763; Gm8764; Gm21304; Gm4340 | 0.54 | 9.03 |
| | | Gm12800 | 0.56 | 8.86 |
| | | Usp17le | 0.84 | 8.83 |
| | | Gm6346 | 0.85 | 8.82 |
| | | H19; Mir675 | 0.92 | 8.56 |
| | | Gm6351 | 0.69 | 8.22 |
| | | Gm7982 | 0.69 | 8.22 |
| | | Gm6509 | 0.84 | 8.16 |
| | | Ly6a | 0.75 | 8.11 |
| | | Gm11543 | 0.59 | 7.73 |
| | | Gm7682 | 0.74 | 7.62 |
| | | Gm5662 | 0.57 | 7.32 |
| | | Gm12794 | 0.43 | 7.30 |
| | | Plet1 | 0.30 | 6.64 |
| | | Gm21818 | 1.20 | 6.41 |

| | | | | |
|---|---|---|---|---|
| | | Gm13128 | 0.19 | 6.30 |
| | | Gm7942 | 0.95 | 6.09 |
| | | Gm2022 | 0.77 | 6.05 |
| | | Got1l1 | 0.91 | 5.95 |
| | | Scml2 | 0.70 | 5.54 |
| | | Gm7963 | 0.81 | 5.42 |
| | | Aqp3 | 1.35 | 5.40 |
| | | Pramel3 | 0.84 | 5.31 |
| | | Gm2808; Tulp4 | 0.54 | 5.27 |
| | | Rad21l | 1.85 | 5.09 |
| | | Eif2s3y | 0.13 | 5.08 |
| | | Sytl3 | 0.51 | 5.05 |
| | | Tdpoz3 | 1.17 | 4.89 |
| | | Gm5286 | 0.77 | 4.87 |
| | | Gm4222 | 0.90 | 4.86 |
| | | Gm8300 | 0.62 | 4.80 |
| | | Usp17la | 0.81 | 4.71 |
| | | Cd80 | 0.89 | 4.70 |
| | | Plekhg1 | 0.82 | 4.67 |
| | | Arl14epl | 0.43 | 4.61 |
| | | Timm8a2 | 0.34 | 4.59 |
| | | Uba1y | 0.50 | 4.54 |
| | | Xlr3c | 0.74 | 4.46 |
| | | Cyr61 | 1.06 | 4.35 |
| | | Gm38423; Gm11239; Gm11236; Gm11237 | 0.95 | 4.29 |
| | | Gm15023; AV320801 | 0.83 | 4.27 |
| | | 4930591A17Rik | 0.83 | 4.25 |
| | | Gm11237; Gm11236 | 0.93 | 4.20 |
| | | Inpp5d | 0.37 | 4.14 |
| | | Gm428 | 0.92 | 4.09 |
| | | Xlr3d-ps | 0.81 | 4.06 |
| | | Gm4778 | 0.90 | 4.02 |
| | | LOC106029237 | 0.70 | 4.01 |
| | | Obox4-ps38 | 0.86 | 3.91 |
| | | Cacnb3 | 0.70 | 3.89 |
| | | Gm11238 | 0.80 | 3.87 |
| | | Dusp9 | 0.76 | 3.67 |
| | | Gm20433; RP24-89F23.1 | 1.31 | 3.64 |
| | | Gadd45g | 1.39 | 3.58 |
| | | Ddx3y | 0.05 | 3.56 |
| | | Gm13119 | 1.09 | 3.53 |
| | | Nlrp9b | 0.78 | 3.52 |
| | | Klhl13 | 0.25 | 3.51 |
| | | Pramef17 | 0.20 | 3.51 |
| | | BB287469 | 0.53 | 3.49 |
| | | Uty | 0.16 | 3.48 |
| | | Gm10663 | 1.40 | 3.47 |
| | | Naprt | 0.99 | 3.46 |
| | | Gm7903; Gm5128; Pramel3 | 0.90 | 3.43 |
| | | LOC100861811 | 0.80 | 3.39 |
| | | E530001F21Rik | 0.69 | 3.38 |
| | | Mep1b | 0.97 | 3.37 |

| | | | | |
|---|---|---|---|---|
| | | 1700013H16Rik | 0.89 | 3.34 |
| | | Gm22862 | 1.04 | 3.34 |
| | | Gm4027; BB287469 | 0.56 | 3.31 |
| | | LOC100861827 | 0.86 | 3.31 |
| | | Gm10944 | 0.73 | 3.30 |
| | | Calcoco2 | 0.24 | 3.26 |
| | | Grasp | 1.12 | 3.25 |
| | | Capn3 | 0.66 | 3.23 |
| | | LOC545763 | 0.84 | 3.23 |
| | | Magea4 | 0.89 | 3.23 |
| | | Nckap1l | 0.55 | 3.22 |
| | | Slc23a1 | 0.46 | 3.20 |
| | | Tbrg3 | 1.35 | 3.20 |
| | | Tnfrsf8 | 0.99 | 3.20 |
| | | Gm16429 | 0.86 | 3.19 |
| | | Gm7971 | 0.86 | 3.19 |
| | | 3830417A13Rik | 0.84 | 3.17 |
| | | 2900011O08Rik | 1.03 | 3.15 |
| | | Ms4a10 | 1.05 | 3.15 |
| | | Slc25a20 | 0.93 | 3.07 |
| | | Usp17lc | 0.78 | 3.07 |
| | | Gm11237; Gm11236; Gm11239 | 0.90 | 3.02 |
| | | Ighv1-62-2 | 0.78 | 2.99 |
| | | Ighv1-71 | 0.78 | 2.99 |
| | | 1700007K13Rik | 1.80 | 2.97 |
| | | Rassf9 | 0.93 | 2.97 |
| | | Gm16203; AC124198.1 | 0.79 | 2.95 |
| | | Mylpf | 0.52 | 2.92 |
| | | Glod5 | 1.04 | 2.89 |
| | | Gm15097 | 1.20 | 2.89 |
| | | Xlr3e-ps | 0.95 | 2.84 |
| | | Mfap5 | 0.88 | 2.83 |
| | | Gm15127 | 1.27 | 2.81 |
| | | Gm7903 | 0.95 | 2.81 |
| | | Tmem92 | 0.77 | 2.81 |
| | | Sfmbt2 | 0.72 | 2.79 |
| | | Gm10424 | 0.86 | 2.78 |
| | | Stmn2 | 1.09 | 2.77 |
| | | Myo10 | 0.83 | 2.76 |
| | | Peg3 | 1.33 | 2.76 |
| | | Gm15698 | 0.47 | 2.71 |
| | | Pramel6 | 1.03 | 2.69 |
| | | Anxa8 | 1.07 | 2.68 |
| | | Glipr2 | 0.85 | 2.67 |
| | | Pdzd2 | 0.80 | 2.65 |
| | | Xlr3b | 0.85 | 2.64 |
| | | Mmrn2 | 0.94 | 2.63 |
| | | Gm14569 | 0.79 | 2.61 |
| | | Xlr3a | 0.90 | 2.61 |
| | | AA623943 | 0.84 | 2.58 |
| | | Slc5a4b | 0.78 | 2.58 |
| | | Taf9b | 0.96 | 2.58 |
| | | Gm20767 | 0.95 | 2.55 |

| | | | | |
|---|---|---|---|---|
| | | Cox7b2 | 0.65 | 2.53 |
| | | Gtse1 | 1.07 | 2.53 |
| | | Impact | 0.66 | 2.52 |
| | | Xlr5c | 0.91 | 2.50 |
| | | Gm2046 | 0.62 | 2.49 |
| | | Nup62cl | 0.90 | 2.49 |
| | | C130073F10Rik | 0.99 | 2.47 |
| | | Gm11546 | 0.70 | 2.47 |
| | | Arhgap8 | 0.54 | 2.46 |
| | | Otud6a | 1.16 | 2.46 |
| | | Gm15093; Gm15085; Ott | 1.32 | 2.45 |
| | | Gm15085; Gm15080; Ott | 1.27 | 2.44 |
| | | Gm11545 | 0.97 | 2.43 |
| | | Pacsin2 | 1.02 | 2.42 |
| | | Zfp706 | 0.77 | 2.42 |
| | | Dusp27 | 0.65 | 2.41 |
| | | Cd55 | 0.82 | 2.39 |
| | | Snord43 | 0.69 | 2.39 |
| | | Gm2035 | 0.63 | 2.38 |
| | | Ifitm3 | 0.24 | 2.38 |
| | | Mbnl2 | 0.84 | 2.38 |
| | | Alox5ap | 0.50 | 2.37 |
| | | BC021614 | 0.90 | 2.37 |
| | | Gm11800 | 0.91 | 2.37 |
| | | Gm15091 | 1.16 | 2.37 |
| | | Tsx | 0.90 | 2.37 |
| | | Gm8994 | 0.89 | 2.36 |
| | | LOC666331; Gm11635; RP23-382C19.4 | 1.07 | 2.36 |
| | | Gm15107; Gm15114; Gm15093; Gm15085; Ott | 1.30 | 2.35 |
| | | Eno3 | 1.29 | 2.34 |
| | | Gm6803 | 0.93 | 2.34 |
| | | Gprc5a | 0.60 | 2.34 |
| | | Gm15128; Gm15107 | 1.25 | 2.32 |
| | | Gm10193 | 0.69 | 2.31 |
| | | Kcnq1ot1 | 1.00 | 2.31 |
| | | Xlr5b | 1.00 | 2.30 |
| | | Gm2075 | 0.97 | 2.28 |
| | | Pop1 | 0.96 | 2.28 |
| | | Steap3 | 0.84 | 2.28 |
| | | Luzp4 | 1.17 | 2.27 |
| | | Mlana | 0.89 | 2.27 |
| | | Dpp4 | 0.83 | 2.26 |
| | | Snora34; Mir1291 | 1.69 | 2.25 |
| | | Gm5890 | 1.06 | 2.22 |
| | | Hip1 | 1.40 | 2.22 |
| | | LOC100861819 | 0.78 | 2.22 |
| | | AI481877 | 0.85 | 2.20 |
| | | Itga5 | 0.72 | 2.20 |
| | | Pramef12 | 0.57 | 2.19 |
| | | Zrsr1 | 1.11 | 2.19 |
| | | Pdxp | 1.85 | 2.18 |

| | | | | |
|---|---|---|---|---|
| | | Plin3 | 1.04 | 2.18 |
| | | 1110038F14Rik | 0.85 | 2.17 |
| | | Elf3 | 0.99 | 2.17 |
| | | Wbscr27 | 0.85 | 2.17 |
| | | Cpne5 | 1.01 | 2.16 |
| | | Gabarapl2 | 0.92 | 2.16 |
| | | Gm11622 | 1.40 | 2.16 |
| | | Prps2 | 1.60 | 2.16 |
| | | Sept1 | 0.09 | 2.16 |
| | | Acad11 | 1.12 | 2.15 |
| | | B4galt6 | 0.78 | 2.15 |
| | | Foxi3 | 0.86 | 2.14 |
| | | Adrb3 | 0.66 | 2.13 |
| | | Gm8332 | 0.94 | 2.13 |
| | | Plk3 | 0.71 | 2.13 |
| | | Cd37 | 1.16 | 2.12 |
| | | Gm10439; Gm15093; Gm15085; Ott | 1.20 | 2.12 |
| | | Olfr836 | 0.86 | 2.12 |
| | | Rad1 | 0.97 | 2.12 |
| | | Gm2056 | 0.93 | 2.11 |
| | | Rasgrp3 | 0.86 | 2.11 |
| | | Slc25a17 | 1.09 | 2.11 |
| | | Tars | 0.70 | 2.10 |
| | | Gm7969 | 0.99 | 2.09 |
| | | Mfge8 | 1.26 | 2.09 |
| | | Oas1a | 0.68 | 2.09 |
| | | Usp26 | 0.20 | 2.09 |
| | | Arih2 | 0.99 | 2.08 |
| | | Gm25956 | 1.30 | 2.08 |
| | | Gm4303; Gm4305; Gm4307; Gm4302 | 0.93 | 2.08 |
| | | Slc7a1 | 0.75 | 2.08 |
| | | Adamts14 | 1.13 | 2.07 |
| | | Car2 | 1.38 | 2.07 |
| | | Morc1 | 0.78 | 2.07 |
| | | Tbc1d22a | 1.03 | 2.07 |
| | | Tmem181b-ps; Tmem181a | 1.56 | 2.07 |
| | | Atg14 | 0.99 | 2.06 |
| | | Fth-ps2 | 1.33 | 2.06 |
| | | Maf1 | 1.10 | 2.06 |
| | | Xlr5a | 0.86 | 2.06 |
| | | Abcb1a | 0.64 | 2.05 |
| | | Smyd1 | 0.76 | 2.05 |
| | | Abcb1b | 0.75 | 2.04 |
| | | Alg14 | 0.80 | 2.04 |
| | | Ctgf | 0.51 | 2.04 |
| | | Gm12790 | 0.67 | 2.04 |
| | | Klk1 | 0.75 | 2.04 |
| | | Usp17ld | 0.87 | 2.04 |
| | | Gm19486 | 1.05 | 2.03 |
| | | Cdkn1a | 0.99 | 2.02 |
| | | Ddit4 | 1.46 | 2.02 |
| | | Obox4-ps26 | 0.76 | 2.02 |
| | | Camk1d | 0.58 | 2.01 |

| | | | | |
|---|---|---|---|---|
| | | Gm21936 | 0.85 | 2.01 |
| | | Gm8701 | 0.90 | 2.01 |
| | | Pim3 | 0.65 | 2.01 |
| | | Bop1 | 0.72 | 2.00 |
| | | Slc38a2 | 0.93 | 2.00 |

### Table S4

#### List of down-regulated gnes in ΔbHLHZ and/or ΔT ESCs

Fold change

>50

0.1>

| | | Δ bHLHZ | ΔT |
|---|---|---|---|
| ΔHLHZ ≦ 0.5 ; ΔT > 0.5 | Ddx3y | 0.05 | 3.56 |
| | Sept1 | 0.09 | 2.16 |
| | Eif2s3y | 0.13 | 5.10 |
| | Fmr1nb | 0.14 | 0.84 |
| | Uty | 0.16 | 3.48 |
| | Zscan4b | 0.19 | 19.70 |
| | Gm13128 | 0.19 | 6.32 |
| | Usp26 | 0.20 | 2.10 |
| | Gm11096 | 0.20 | 0.88 |
| | Pramef17 | 0.20 | 3.51 |
| | Aim2 | 0.21 | 1.13 |
| | Zp3 | 0.21 | 1.39 |
| | Zscan4f | 0.22 | 15.14 |
| | Hmgn5 | 0.22 | 0.81 |
| | AU018091 | 0.23 | 1.54 |
| | Zscan4a | 0.23 | 17.27 |
| | Rhox5 | 0.24 | 18.64 |
| | Calcoco2 | 0.24 | 3.25 |
| | Ifitm3 | 0.24 | 2.38 |
| | Fam111a | 0.25 | 0.52 |
| | Naa11 | 0.25 | 0.73 |
| | Klhl13 | 0.25 | 3.51 |
| | Lgals9 | 0.26 | 0.95 |
| | Smoc1 | 0.26 | 1.40 |
| | Zscan4e | 0.27 | 19.84 |
| | Fbxo15 | 0.27 | 1.80 |
| | Mir1950 | 0.29 | 0.72 |
| | CN725425 | 0.29 | 1.23 |
| | Plet1 | 0.30 | 6.63 |
| | Id2 | 0.30 | 0.92 |
| | Zscan4c | 0.30 | 15.89 |
| | Zscan4d | 0.31 | 13.55 |
| | Ulbp1 | 0.33 | 0.63 |
| | Coch | 0.34 | 0.60 |
| | Timm8a2 | 0.34 | 4.59 |
| | Lrrc2 | 0.35 | 1.83 |
| | Elmo1 | 0.35 | 0.97 |
| | 2410137M14Rik | 0.36 | 1.85 |
| | Hist1h2aa | 0.36 | 5.46 |
| | Inpp5d | 0.37 | 4.14 |

| | | Δ bHLHZ | ΔT |
|---|---|---|---|
| ΔHLHZ & ΔT ≦ 0.5 | Gm11114 | 0.14 | 0.25 |
| | Ighv1-12 | 0.30 | 0.21 |
| | Krt17 | 0.33 | 0.39 |
| | Gm21028 | 0.34 | 0.36 |
| | P3h4 | 0.36 | 0.49 |
| | Gm21954 | 0.37 | 0.37 |
| | Magohb | 0.41 | 0.40 |
| | Mboat1 | 0.41 | 0.44 |
| | Pmel | 0.41 | 0.20 |
| | Babam1 | 0.43 | 0.44 |
| | Spink1 | 0.50 | 0.24 |
| ΔT ≦ 0.5 ; ΔHLHZ > 0.5 | Gm15428 | 0.77 | 0.14 |
| | Phlda2 | 0.75 | 0.16 |
| | Lefty2 | 1.37 | 0.19 |
| | Gm10087 | 0.91 | 0.19 |
| | Ttll4 | 1.06 | 0.23 |
| | Grb10 | 1.09 | 0.23 |
| | 2810417H1 | 0.92 | 0.25 |
| | Ap1m2 | 0.71 | 0.27 |
| | Gm11115 | 1.06 | 0.28 |
| | Spred1 | 0.80 | 0.30 |
| | Rpusd2 | 1.21 | 0.32 |
| | Arxes2 | 0.71 | 0.32 |
| | Fgf15 | 1.38 | 0.33 |
| | Stx3 | 1.30 | 0.32 |
| | Rragb | 1.25 | 0.33 |
| | Tmem184c | 0.67 | 0.33 |
| | Cldn6 | 0.78 | 0.34 |
| | Gm2862 | 0.92 | 0.34 |
| | Nefl | 1.10 | 0.34 |
| | Tmem47 | 1.19 | 0.34 |
| | Hook1 | 0.61 | 0.35 |
| | Ly75 | 1.21 | 0.35 |
| | Wdr83os | 0.58 | 0.36 |
| | Rtf1 | 0.86 | 0.36 |
| | Fgf17 | 0.81 | 0.37 |
| | Zfp280c | 1.09 | 0.37 |
| | Cdh1 | 0.73 | 0.37 |
| | Slain1 | 0.89 | 0.37 |
| | Egr1 | 1.08 | 0.38 |

| | | | |
|---|---|---|---|
| | Rlbp1 | 0.37 | 1.20 |
| | Cdh9 | 0.37 | 1.35 |
| | Id1 | 0.37 | 1.02 |
| | Capsl | 0.37 | 1.16 |
| | Tmx3 | 0.38 | 0.77 |
| | Laptm4b | 0.38 | 0.52 |
| | Tspan6 | 0.39 | 1.00 |
| | Liph | 0.39 | 3.61 |
| | Sohlh2 | 0.39 | 1.39 |
| | Olfr985 | 0.40 | 1.69 |
| | Mme | 0.40 | 1.62 |
| | Gm11489 | 0.40 | 0.66 |
| | Thoc5 | 0.41 | 0.87 |
| | Vamp7 | 0.41 | 0.99 |
| | S100a11 | 0.41 | 1.00 |
| | Gab1 | 0.41 | 0.66 |
| | Klf12 | 0.41 | 1.00 |
| | 9430069I07Rik | 0.41 | 1.38 |
| | Fancl | 0.41 | 0.61 |
| | Fam25c | 0.42 | 0.69 |
| | H2-M6-ps | 0.42 | 1.69 |
| | Folr1 | 0.42 | 1.44 |
| | Mgmt | 0.43 | 1.57 |
| | Arl14epl | 0.43 | 4.59 |
| | Gpx2-ps1 | 0.43 | 0.78 |
| | Gm12794 | 0.43 | 7.31 |
| | Anxa2 | 0.43 | 0.82 |
| | Dmc1 | 0.43 | 1.55 |
| | Gm9631 | 0.44 | 0.53 |
| | Tek | 0.45 | 1.07 |
| | Slc18b1 | 0.45 | 1.33 |
| | Jam2 | 0.45 | 1.42 |
| | Atf4 | 0.45 | 1.56 |
| | Gys1 | 0.45 | 0.82 |
| | Trap1a | 0.45 | 0.82 |
| | Slc23a1 | 0.46 | 3.20 |
| | Slc25a12 | 0.46 | 1.32 |
| | Psap | 0.47 | 2.28 |
| | Nxf3 | 0.47 | 1.04 |
| | Cth | 0.47 | 1.39 |
| | Gm11575 | 0.47 | 0.67 |
| | Golph3l | 0.47 | 0.81 |
| | Gm20939 | 0.47 | 0.61 |
| | H2-M5 | 0.47 | 1.09 |

| | | | |
|---|---|---|---|
| | Gm2573 | 0.69 | 0.38 |
| | Arxes1 | 0.70 | 0.39 |
| | Spint1 | 1.17 | 0.39 |
| | Bcl6b | 0.82 | 0.39 |
| | Scd1 | 1.13 | 0.39 |
| | Bub1b | 1.26 | 0.40 |
| | Gm12245 | 0.77 | 0.40 |
| | Ifi30 | 1.42 | 0.41 |
| | Lefty1 | 1.39 | 0.42 |
| | Oxa1l | 1.60 | 0.43 |
| | Nusap1 | 1.44 | 0.43 |
| | Tmem203 | 0.60 | 0.43 |
| | Casc5 | 1.02 | 0.43 |
| | Mir669b | 0.74 | 0.43 |
| | Hook2 | 0.98 | 0.43 |
| | Ighv4-1 | 0.63 | 0.43 |
| | 4931400O( | 0.62 | 0.44 |
| | Smim13 | 0.77 | 0.44 |
| | Gm19505 | 1.09 | 0.44 |
| | Rasgrp1 | 0.76 | 0.44 |
| | Ino80 | 0.74 | 0.44 |
| | Ptpn3 | 1.04 | 0.44 |
| | Gm20721 | 1.27 | 0.44 |
| | Ier2 | 0.93 | 0.44 |
| | Rpap1 | 1.09 | 0.44 |
| | Rad51 | 1.01 | 0.45 |
| | Zfp819 | 0.59 | 0.45 |
| | Gm12255 | 0.77 | 0.45 |
| | Zfp953; ZF | 1.00 | 0.45 |
| | Shcbp1 | 0.73 | 0.45 |
| | Ndrg2 | 1.45 | 0.45 |
| | Zfp459 | 1.09 | 0.46 |
| | Trh | 0.85 | 0.46 |
| | Tecr | 0.70 | 0.46 |
| | Rnaseh2b | 1.57 | 0.46 |
| | Bspry | 0.90 | 0.46 |
| | Rbmxl1 | 0.57 | 0.47 |
| | Oip5 | 1.04 | 0.47 |
| | Gsr | 0.75 | 0.47 |
| | Cdh3 | 1.05 | 0.47 |
| | Asna1 | 0.73 | 0.47 |
| | Aldoc | 1.22 | 0.47 |
| | Rcbtb1 | 1.93 | 0.47 |
| | Isoc1 | 0.68 | 0.47 |

| | | | |
|---|---|---|---|
| | Gm15698 | 0.47 | 2.71 |
| | Lsm8; Naa38 | 0.48 | 0.81 |
| | Pcolce2 | 0.48 | 0.65 |
| | Hck | 0.48 | 1.26 |
| | Prpsap1 | 0.48 | 0.99 |
| | Pof1b | 0.48 | 0.55 |
| | Efhd1 | 0.48 | 0.65 |
| | Lipa | 0.48 | 0.57 |
| | Pstpip2 | 0.48 | 0.74 |
| | Ighv2-2 | 0.48 | 0.68 |
| | Fkbp10 | 0.48 | 0.52 |
| | Mrpl34 | 0.48 | 0.78 |
| | Chmp2a | 0.49 | 1.87 |
| | Gm7889 | 0.49 | 1.24 |
| | Ddx49 | 0.49 | 0.56 |
| | Gm10177 | 0.49 | 0.81 |
| | Casp3 | 0.49 | 0.54 |
| | D1Pas1 | 0.49 | 4.35 |
| | 1810037I17Rik | 0.49 | 0.84 |
| | Pla1a | 0.49 | 1.74 |
| | Mpv17l2 | 0.49 | 0.70 |
| | Hmox1 | 0.49 | 1.03 |
| | As3mt | 0.49 | 1.31 |
| | Gm21060 | 0.49 | 0.62 |
| | Pnp | 0.49 | 0.97 |
| | Cdkn1c | 0.49 | 1.07 |
| | Lrrc31 | 0.49 | 0.88 |
| | Msantd4 | 0.49 | 0.61 |
| | Eci2 | 0.50 | 0.95 |
| | Nmnat2 | 0.50 | 1.56 |
| | Ooep | 0.50 | 0.88 |
| | Gm10495 | 0.50 | 0.86 |
| | Uba1y | 0.50 | 3.78 |
| | Alox5ap | 0.50 | 2.38 |

| | | | |
|---|---|---|---|
| | Serpinb6c | 0.57 | 0.47 |
| | Fam98b | 0.90 | 0.47 |
| | Gpc3 | 0.99 | 0.48 |
| | Elovl7 | 0.84 | 0.48 |
| | Arl2bp | 0.55 | 0.48 |
| | Slc25a4 | 0.72 | 0.48 |
| | Supt16 | 2.01 | 0.48 |
| | Vegfb | 0.94 | 0.48 |
| | BC052040 | 1.11 | 0.48 |
| | Dph6 | 1.11 | 0.48 |
| | Hmces | 0.95 | 0.48 |
| | Spint2 | 0.86 | 0.48 |
| | 4921524J1 | 0.53 | 0.48 |
| | Cfdp1 | 0.52 | 0.48 |
| | Nufip1 | 1.52 | 0.48 |
| | Aqr | 0.93 | 0.49 |
| | Thap11 | 0.55 | 0.49 |
| | Hap1 | 0.81 | 0.49 |
| | Rnaseh2a | 0.64 | 0.49 |
| | Mir467c | 1.04 | 0.49 |
| | Smpdl3b | 0.74 | 0.49 |
| | Cers4 | 0.81 | 0.49 |
| | Asf1b | 1.10 | 0.49 |
| | 2310036O2 | 0.60 | 0.49 |
| | Nutf2; Nut | 0.62 | 0.49 |
| | Gng3 | 1.39 | 0.49 |
| | Ccdc115 | 1.23 | 0.50 |
| | Tyro3 | 0.95 | 0.50 |
| | Psmb10 | 0.84 | 0.50 |
| | LOC10263 | 0.82 | 0.50 |
| | Gm8775 | 0.84 | 0.50 |
| | Syce2 | 1.25 | 0.50 |
| | Cd276 | 1.34 | 0.50 |
| | Nab2 | 0.70 | 0.50 |
| | Timm21 | 0.65 | 0.50 |
